# Mechanosensitive channel MscS is critical for termination of the bacterial hypoosmotic permeability response

**DOI:** 10.1101/2023.02.27.530336

**Authors:** Elissa Moller, Madolyn Britt, Anthony Schams, Hannah Cetuk, Andriy Anishkin, Sergei Sukharev

## Abstract

Free-living microorganisms are subjected to drastic changes in osmolarity. To avoid lysis under sudden osmotic down-shock, bacteria quickly expel small metabolites through the tension-activated channels MscL, MscS, and MscK. We examined five chromosomal knockout strains, Δ*mscL*, Δ*mscS*, a double knockout Δ*mscS* Δ*mscK*, and a triple knockout Δ*mscL* Δ*mscS* Δ*mscK* in comparison to the wild-type parental strain. Stopped-flow experiments confirmed that both MscS and MscL mediate fast osmolyte release and curb cell swelling, but osmotic viability assays indicated that they are not equivalent. MscS alone was capable of rescuing the cell population, but in some strains MscL did not rescue and additionally became toxic in the absence of both MscS and MscK. Furthermore, MscS was upregulated in the Δ*mscL* strain, suggesting either a cross-talk between the two genes/proteins or the influence of cell mechanics on *mscS* expression. The data shows that for the proper termination of the permeability response, the high-threshold (MscL) and the low-threshold (MscS/MscK) channels must act sequentially. In the absence of low-threshold channels, at the end of the release phase, MscL should stabilize membrane tension at around 10 mN/m. Patch-clamp protocols emulating the tension changes during the release phase indicated that the non-inactivating MscL, residing at its own tension threshold, flickers and produces a protracted leakage. The MscS/MscK population, when present, stays open at this stage to reduce tension below the MscL threshold and silence the large channel. When MscS reaches its own threshold, it inactivates and thus ensures proper termination of the hypoosmotic permeability response. This functional interplay between the high- and low-threshold channels is further supported by the compromised osmotic survival of bacteria expressing non-inactivating MscS mutants.

**Summary (for the table of contents):** The kinetics of hypotonic osmolyte release from *E. coli* is analyzed in conjunction with bacterial survival. It is shown that MscL, the high-threshold ‘emergency release valve’, rescues bacteria from down-shocks only in the presence of MscS, MscK or other low-threshold channels that are necessary to pacify MscL at the end of the release phase.

## Introduction

Free living bacteria and environmentally transmitted pathogens are regularly subjected to drastic osmolarity changes. Under high-osmolarity conditions they accumulate large amounts of compatible osmolytes such as betaine, proline, and glutamate (Wood et al., 2001). Because bacteria rely on turgor pressure to maintain shape and volume, they accumulate these intracellular osmolytes in concentrations that always exceed the external osmolarity by 200-400 mOsm. In the event of a drastic drop of external osmolarity, such as a sudden rain storm, bacteria quickly release most of their small intracellular osmolytes to avoid lysis. Mechanosensitive channels MscS and MscL, which are directly activated by tension in the cytoplasmic membrane (Kung et al., 2010; Moe and Blount, 2005; Nomura et al., 2012; Sukharev, 2002; Sukharev et al., 1993), were identified as the main components of the osmolyte release system in *E. coli* (Levina et al., 1999). Five additional channels from the MscS family (MscK, YnaI, YbdG, YbiO, and YjeP) were identified in the *E. coli* genome as well but were found to be less abundant under standard laboratory conditions (Edwards et al., 2012). Among them, YbdG (Schumann et al., 2010) and YjeP (Edwards et al., 2012) were associated with sparsely observed mini-channel (MscM) activity. Like *E. coli*, it is common for most bacterial species to have this arrangement comprising two types of channels: the MscL family, with a large conductance and a high activation tension, and the MscS family, with a smaller conductance and a lower activation tension (Balleza and Gomez-Lagunas, 2009). Genomes of other bacteria studied by electrophysiological techniques such as *Vibrio cholerae* (Rowe et al., 2013) or *Pseudomonas aeruginosa* (Cetiner et al., 2017) have one *mscL*, one or two *mscS* genes, and a set of 3-6 more distant MscS homologs whose functions are yet to be identified.

The first description of single and double knockout phenotypes (Levina et al., 1999) suggested that MscS and MscL channels are qualitatively similar in their rescuing ability and appeared partially redundant. However, further detailed functional patch-clamp characterization of excised spheroplast patches and liposomes has shown these channels to be distinct. The high-threshold MscL channel activates at near-lytic tensions 12-14 mN/m (midpoint) by opening an almost non-selective 3 nS pore (Yang and Blount, 2011), acting as a last resort emergency release valve (Belyy et al., 2010; Chiang et al., 2004). Analysis of gating patterns concluded that the channel essentially behaves as a two-state system, transitioning between closed and open states. Its large expansion area (~20 nm^2^) produces a steep response to tension above the threshold, and the channel shows only a slight adaptation, apparently due to mechanical patch relaxation, but no inactivation (Belyy et al., 2010). The low-threshold MscS, on the other hand, activates at sub-lytic tensions of 5-7 mN/m, opening a ~1 nS pore. In contrast to MscL, the small-conductance MscS channel readily and fully activates in response to abrupt tension pulses, but only a fraction of the channel population opens in response to slow tension ramps (Akitake et al., 2005; Belyy et al., 2010; Boer et al., 2011). This behavior was attributed to gradual inactivation of channels from the resting state without opening, which occurs under moderate tensions (Cetiner et al., 2018). Furthermore, inactivation under near-threshold tension was shown to be strongly augmented by increased macromolecular crowding pressure on the cytoplasmic side (Rowe et al., 2014). In the inactivated state MscS is non-conductive and completely tension-insensitive. Upon tension release, the channel recovers to the resting state within 1-3 s. A number of mutations have been found that either increase the rate of inactivation or abolish it entirely (Akitake et al., 2007; Boer et al., 2011; Koprowski et al., 2011; Rowe et al., 2014).

Patch-clamp experiments also indicate that in real-life conditions the combined MscS and MscL populations should mediate a graded osmolyte release response, activating and de-activating sequentially. The end of release, where the decreasing membrane tension passes the thresholds for MscL and then for MscS, is an especially interesting phase that suggests functional interplay between these two populations that helps optimize the osmolyte release process and ensure the fast and complete resealing of the cytoplasmic membrane after the shock.

In the present work we utilize a set of chromosomal deletion mutants derived from the wild-type Frag1 strain to analyze the contributions of each type of channel to the osmolyte release process and osmotic viability. We provide a detailed analysis of the release kinetics using the stopped-flow light scattering technique and emphasize the differences in the swelling, release, and termination phases between the osmotically protected and unprotected strains. We show that although MscL can mediate fast release, in certain contexts it is unable to rescue bacteria even from moderate shocks alone. Conversely, MscS provides full protection over the entire range of shocks. We then dissect the deactivation process of the MscL and MscS populations near their respective threshold tensions in patch-clamp experiments and show that the ‘leaky’ deactivation of MscL is different from that of the inactivating behavior of MscS. MscS inactivation allows for prompt and complete resealing of the membrane, which manifests as a ‘hard stop’ in the stopped-flow light scattering traces. We discuss the interplay between the two channels in which the presence of the low-threshold MscS drives the tension far below the activation threshold for MscL and thus silences it. Subsequently, when the tension reaches the threshold for MscS, the channel inactivates and properly terminates the immense permeability response. The data also suggests that MscK can functionally substitute for MscS.

## Methods

### Strains

Bacterial strains originated from the I. Booth laboratory (University of Aberdeen, UK) and were provided by Dr. Samantha Miller.

Frag1 is F– *rha, thi, gal, lacZ* (Epstein and Kim, 1971) is the parental strain for the MS channel deletion strains listed below.

MJF367 (Δ*mscL*) is Frag1 Δ*mscL*::*Cm* (Levina et al., 1999).
MJF451 (Δ*mscS*) is Frag1 Δ*yggB* (Levina et al., 1999).
PB113 (Δ*mscS*Δ*mscK*) is MJF429 Δ*recA* (Li et al., 2002), provided by Dr. Paul Blount (UT, Dallas).
MJF429 is Frag1 Δ*yggB*, Δ*mscK::kan* (Levina et al., 1999).
MJF465 (Δ*mscL* Δ*mscS* Δ*mscK*) is Frag1 Δ*mscL::Cm*, Δ*yggB*, Δ*mscK::kan* (Levina et al., 1999).

E. coli MscL wild-type (WT), MscS WT, and MscS with point mutations changing either the glycine at position 168 to aspartic acid (Koprowski et al., 2011; Rowe et al., 2014) or the glycine at position 113 (Akitake et al., 2007) to alanine (G168D and G113A respectively) were expressed from the pB10d vector, a modified pB10b plasmid (Iscla et al., 2008). The plasmid is IPTG inducible with the lac UV5 and lacIq promoters, confers Ampicillin resistance, and includes a C-terminal 6-His tag on the MscS gene. It uses the high-copy-number ColE1/pMB1/pBR322/pUC origin of replication and the T7 bacteriophage ribosome binding site.

### Osmotic survival

Osmotic viability was determined by counting colony forming units (CFU’s) /mL after osmotic shock from 1200 mOsm down to a lower osmolarity media and then normalized to the 1200 mOsm dilution control. Cells were sub-cultured 1:100 from standard overnight cultures in Luria Bertani (LB) media into LB supplemented with NaCl to final osmolarity 1200 mOsm and grown to early logarithmic phase (OD_600_ of ~0.25). Plasmid expressed channels were induced with 1 mM IPTG for the final 30 minutes. Environmental down-shock was simulated by rapidly pipetting 50 uL of growth culture into 5 mL of 1200 mOsm LB (control), 800 mOsm LB, 600 mOsm LB, 400 mOsm LB, 300 mOsm LB, 200 mOsm LB, and 100 mOsm LB. Each sample was incubated at room temperature for 15 minutes then diluted 1:100 into the corresponding shock media. Cells were then plated in duplicate and colonies were manually counted the next morning. Fraction of osmotic survival was determined by normalizing cell counts relative to the un-shocked 1200 mOsm control. Shock experiments were performed independently at least 4 times and the error shown is the standard deviation between these trials. Two-tail two-sample t-tests assuming unequal variances were performed to determine significance. The p-values are reported in Table S1 in the supplement.

### Stopped-flow

Scattering depends on the size of the cell and the ratio of the refractive indices inside and outside the cell (Koch, 1961; Koch et al., 1996). Refractive indexes of media were measured with a Rudolph J357 refractometer. Swelling and release of internal osmolytes reduce light scattering and the Stopped-Flow device measures the time course of this change. Cells were sub-cultured 1:240 from standard LB overnight cultures into LB supplemented with NaCl to final osmolarity 1200 mOsm and grown to early logarithmic phase (OD_600_ of ~0.25). Plasmid expressed channels were induced with 1 mM IPTG for the final 30 minutes. Cells were harvested and resuspended in their supernatant to OD_600_ of ~2.0 to concentrate them. Once concentrated, cells were rapidly mixed with shock media using a Stopped-Flow machine (Bio-Logic SFM-2000). The machine is equipped with two independent motorized syringes, an 81 μl optical mixing chamber, modular spectrophotometer, and a computer for protocol programming and data acquisition. The resulting small-angle light-scattering from the suspension upon rapid mixing was measured with a PMT tube. Light-scattering measured at an absorbance of 600 nm (A600) was collected over a period of 4 seconds, with a 10 ms sampling rate. The mixing ratio in all experiments was 1:10 delivered at the total rate of 8.5 ml/s, to achieve an ~10 ms mixing time. Cells were shocked sequentially into each osmolarity media with five technical replicates per trial and the traces were averaged. Background shock readings of each media mixed with the cell-free supernatant were taken in the same manner and subtracted from cellular shocks to account for any light-scattering effects of the growth or shock media itself and act as a baseline measurement.

To extract the extent of cell swelling and the fraction of osmolyte release we generated ‘calibration’ curves using the full form of Rayleigh-Gans equation (Supplement sections S2 and S3). The determination of the rate of osmolyte release was done by fitting light scattering traces using MATLAB to the simplified form of the Rayleigh-Gans equation 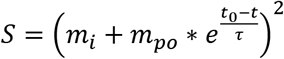 where *m_i_* and *m_po_* are the masses of impermeable and initial mass of permeable osmolytes inside the cell, *t_0_* designates the start of the fit, *t* is time and 1/τ is the rate of osmolyte release. This simplified form was adopted previously (Cetiner et al., 2017). For fitting in all cases, t_0_ was chosen as the time at which the negative slope of the scattering data was steepest, i.e., the minimum of the first derivative, which indicates the point of the fastest release and maximum channel activity. The time to the steepest point was recorded. The fit continued until this slope leveled off, when channels close. The point where the swelling ends and release begins corresponds to the minimum of the second derivative. The time to the release onset was also analyzed. The extent of the swelling is the initial decrease of scattering before the release onset. The fraction of permeable osmolytes is taken from the initial scattering to the point where the traces level off. The final slope was calculated for the final 100 ms of the light scattering trace. At least 4 biological replicates of each strain were performed, the fits were averaged, and the standard deviation was taken between these. All of the knockout strains were down-shocked into 800, 600, 400, 300, 200, and 100 mOsm LB. Plasmid expressed channels were down-shocked into 900, 600, 450, 300, and 150 mOsm LB with a final down-shock into water. 800 and 900 mOsm were not included for extent of swelling, time to release, or slope due to the majority of the traces being comprised purely of swelling without release at these osmolarities.

### Patch-clamp

Giant spheroplasts were generated as described previously (Martinac et al., 1987). Briefly, cells were grown in LB to turbidity and then sub-cultured 1:9 into MLB (250 mOsm) and grown for 1.5 h in the presence of 0.06 mg/mL cephalexin, which prevents cell septation. MscS expressed from plasmids was induced with 1 mM IPTG for the final 30 minutes. The resulting filaments were transferred into 1 M sucrose with 1.8 mg/mL BSA buffer and treated with 0.17 mg/mL lysozyme in the presence of 0.004 M EDTA, which degrades the peptidoglycan layer and causes the filaments to collapse into giant spheres. The reaction was stopped with Mg^2+^. The giant spheroplasts were isolated via sedimentation through a one-step sucrose gradient.

Patch-clamp traces were recorded from giant bacterial cell spheroplasts. Population channel recordings were conducted at +30 mV (pipette) on excised inside-out patches in symmetric buffer containing 400 mM sucrose, 200 mM KCl, 50 mM MgCl_2_, 5 mM CaCl_2_, and 5 mM HEPES at pH 7.4. Membrane patches were obtained using borosilicate glass pipettes (Drummond) and recorded on an Axopatch 200B amplifier (Molecular Devices). Negative pressure stimuli were applied using a high-speed pressure clamp apparatus (HSPC-1; ALA Scientific) programmed using the Clampex software (Molecular Devices). Pressure and voltage protocol programming and data acquisition were performed with the PClamp 10 suite (Axon Instruments). The patch responses were first recorded at +30 mV (pipette) with symmetric 1-s pressure ramps to observe the two waves of current and to determine the activation thresholds, midpoints and saturation levels for the MscS and MscL channel populations. The standard pipettes were pulled to bubble # of 5.3-5.5. Based on the previously reported tension midpoints for both MscS and MscL (Belyy et al., 2010; Moe and Blount, 2005; Nomura et al., 2012; Sukharev, 2002; Sukharev et al., 1999), the radius of a stretched hemispherical patch was estimated as 1.4±0.1 μm. A full table of patch-clamp channel numbers and gating pressures can be found in the supplement in Table S4. To reveal how the channels behave at their tension thresholds, the full population current was activated with a step and then the pressure was ramped down to the threshold of the respective channel. In some experiments, the populations were held at tensions 10 to 30 mm Hg below the respective tension threshold. To observe the inactivation of MscS, a pulse-step-pulse protocol (Akitake et al., 2005; Kamaraju et al., 2011) was used: the full population of channels was opened with a saturating test pulse and then held at the midpoint tension for MscS for 10s followed by a final test pulse again. Percent inactivation was taken from one minus the final pulse normalized to the initial pulse.

### RT-qPCR

The strains Frag1, MJF367 (Δ*mscL*), and MJF451 (Δ*mscS*) were grown in LB to mid log phase (OD_600_ of ~0.25). Total RNA was extracted from cell cultures using the Monarch Total RNA Miniprep Kit (New England Biolabs) according to the manufacturer’s protocol. The concentration of RNA was measured with the NanoDrop OneC (Thermo Fisher Scientific). RNA samples were then reverse transcribed into cDNA using the qScript cDNA Synthesis Kit (Quantabio), resulting in a final concentration of 20 ng/uL. The reverse transcription reaction was carried out at 22 °C for 5 min, 42 °C for 30 min, and 85 °C for 5 min. qRT-PCR was carried out on a ROCHE LC480 using the PowerUp SYBR Green Master Mix (Thermo Fisher Scientific) according to the manufacturer’s protocol. Five technical replicates were run for four biological replicates of each strain with primers specific for *mscL, mscS, mscK*, and *rpmA*. The experimental cycling conditions were 50 °C for 2 minutes then 95 °C for 2 minutes, followed by 40 cycles of 95 °C for 15 seconds, 60 °C for 15 seconds, and 72 °C for 1 min. Data are presented as expression fold change using the 2^-ΔΔCT^ method relative to the housekeeping gene for the 50S ribosomal protein L27 (rpmA) (Ionescu and Belkin, 2009; Livak and Schmittgen, 2001). The error shown in Figure 6B is the standard deviation between replicates. A one-tail two-sample t-test assuming unequal variances was performed to determine significance. Primer efficiency was determined using cDNA from both strains diluted in a 1:10 dilution scheme resulting in concentrations ranging from 1 ng/uL to 0.01 ng/uL with six technical replicates for each primer (Wong and Medrano, 2005). The cycling conditions were 95 °C for 5 minutes followed by 40 cycles of 95 °C for 5 seconds and 60 °C for 30 seconds. A melt curve was run after the cycles for both the experimental conditions and the primer efficiency consisting of 95 °C for 5 seconds, 65 °C for 1 minute, and 97 °C with a ramp rate of 0.11 °C/second. Both the primer efficiency data and the experimental data were analyzed using Abs Quant/ Fit Points to determine the Ct’s and Tm Calling to visualize melt curves and confirm that only one amplicon was present. The averages were taken between the technical replicate Ct’s. Based on these efficiencies and the average Ct’s, the relative expression ratio was calculated using the equation 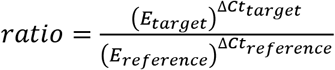 where E is the primer efficiency 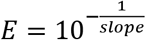. See Supplemental section S5 for the primers and primer efficiency plots.

**Figure 1.**
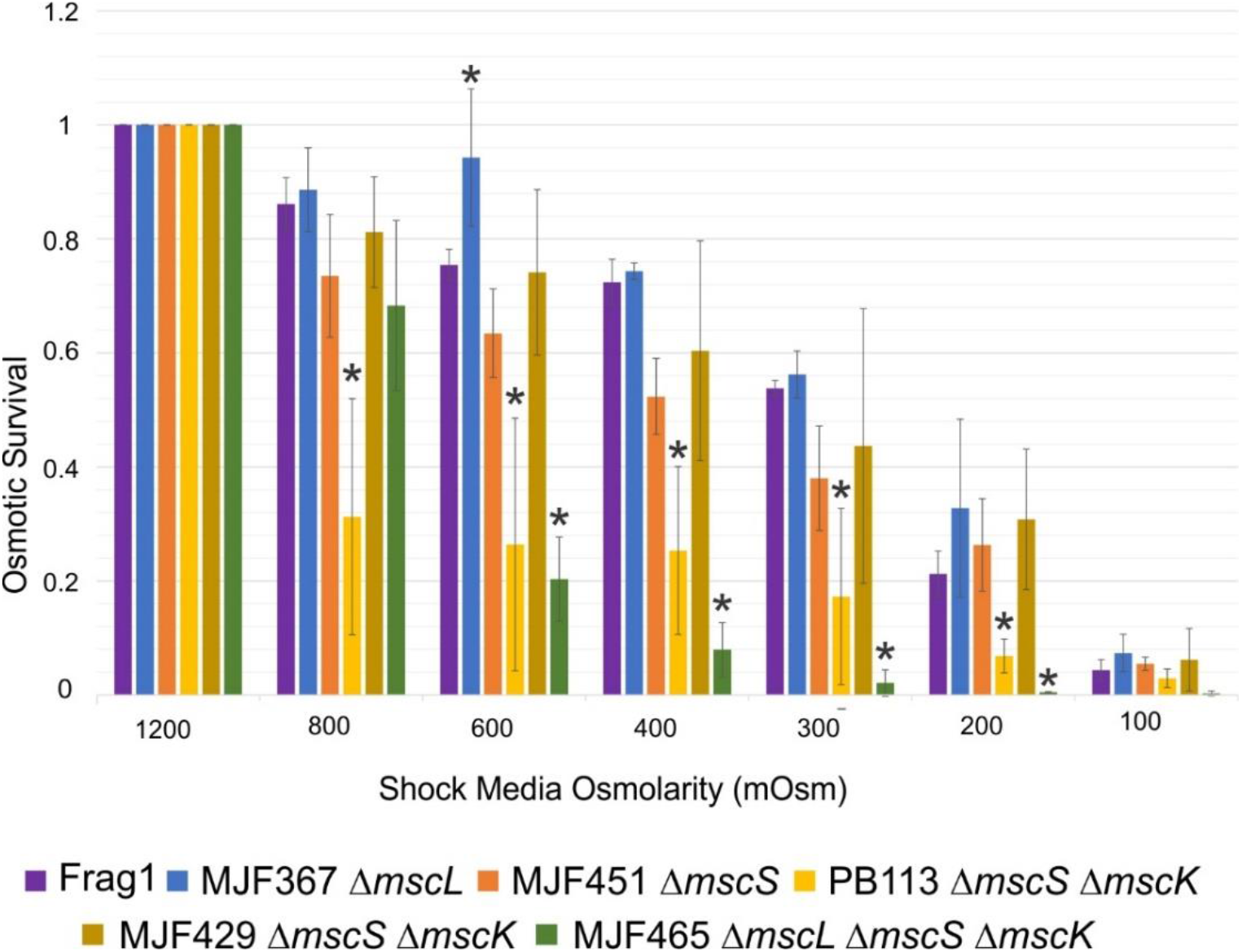
The osmotic viability of the Frag1 parental strain (WT) and the five MS channel deletion strains. Statistically significant (p<0.05) survival values when compared to the Frag1 strain are represented with an asterisk. A full table of all p-values can be found in Supplemental Table S1. PB113 is a *recA-* version of MJF429.

**Figure 2.**
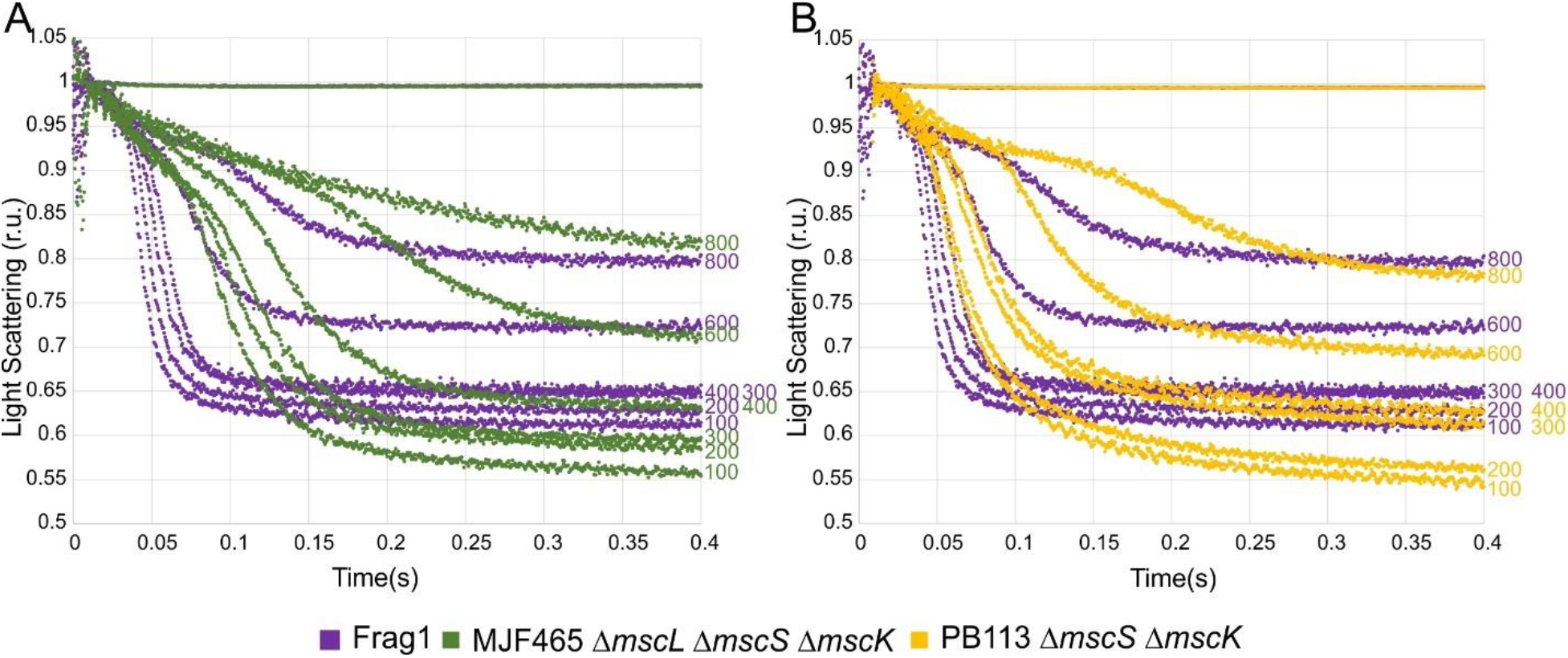
The kinetics of osmolyte release in the triple knockout MJF465 strain (A) is visibly delayed compared to the parent strain Frag1. The kinetics of release in the PB113 strain carrying primarily MscL (B) has considerably shorter characteristic times of osmolyte release, but similar ending to the release from the ‘empty’ MJF465. This continued release, indicated by the creeping end level and the higher fraction of permeable osmolytes, is common for both of these osmotically vulnerable strains (see Figure 1).

**Figure 3.**
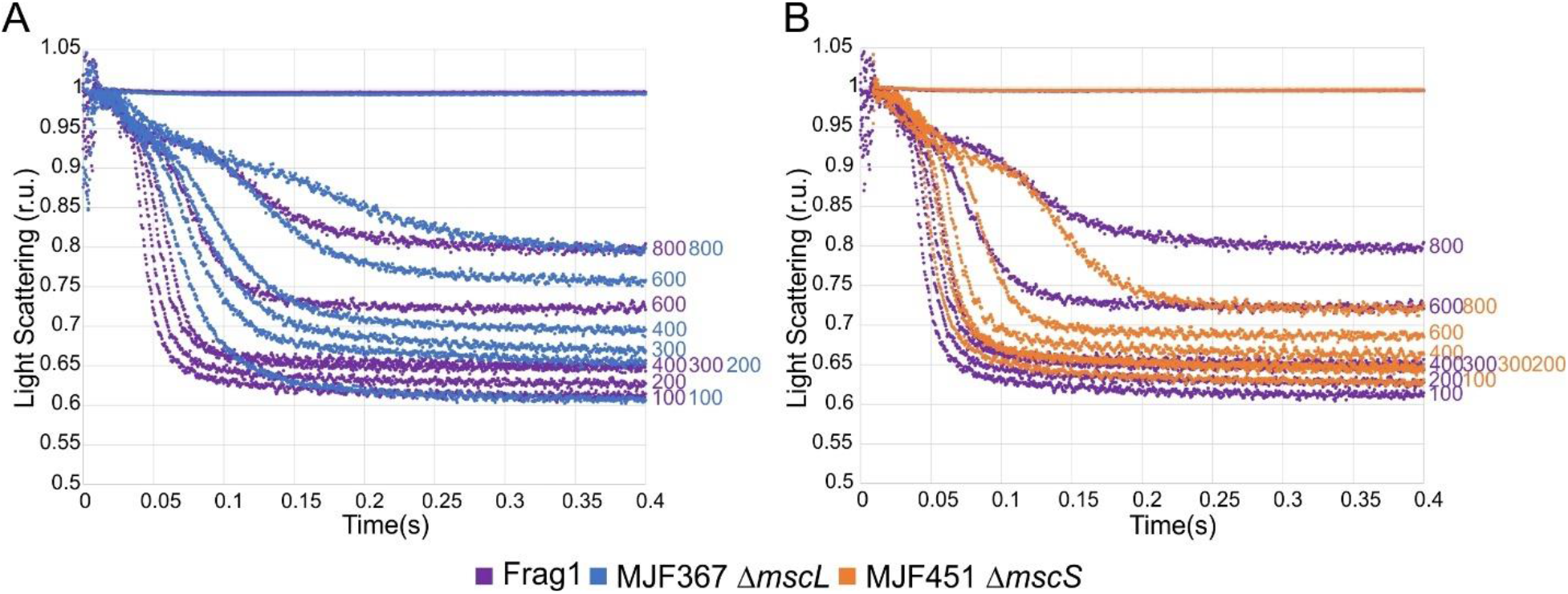
The comparison of osmolyte release kinetics from MJF367 and MJF451 strains. MJF367 shows a slightly extended time of release. Both show a ‘hard stop’ where scattering plateaus after release and survive comparably or better than Frag1 (see Figure 1).

**Figure 4.**
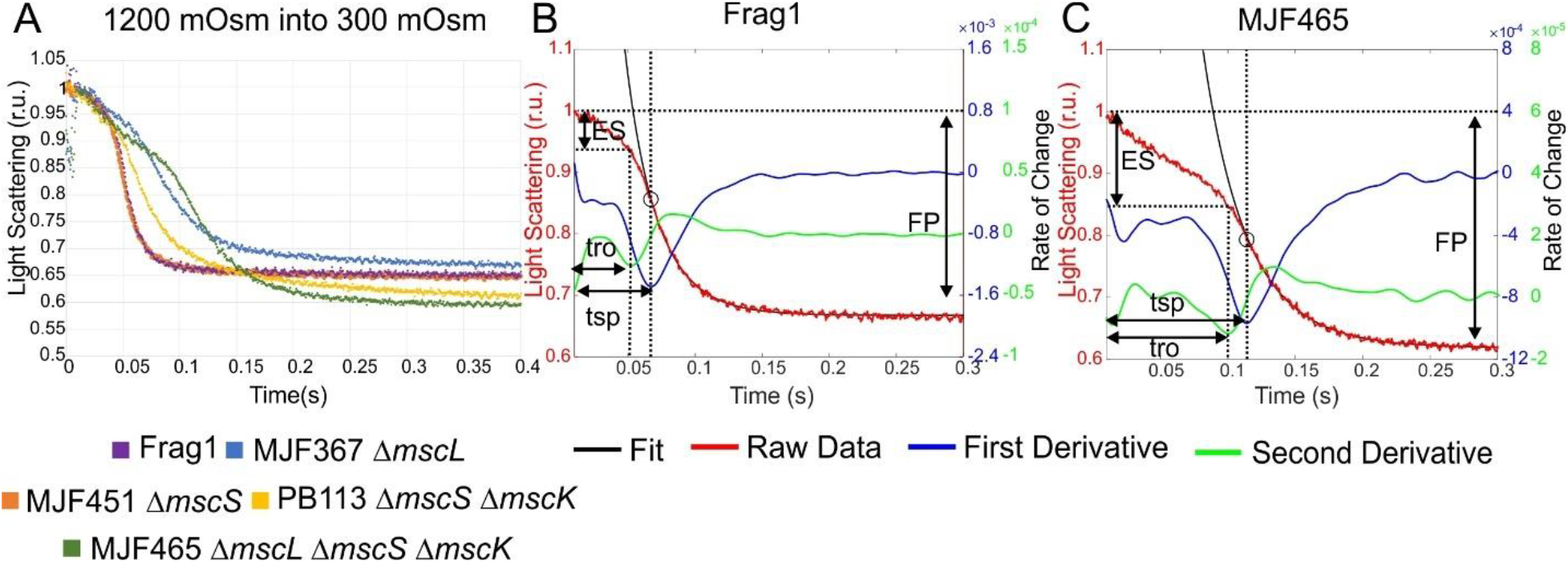
The characteristic stopped-flow traces and their treatment. The light scattering traces after fast mixing from 1200 to 300 mOsm are shown comparatively for all five strains (A). The fitting scheme is shown for Frag1 (B) and MJF465 (C). The experimental traces are shown in red. The minimum of the first derivative (blue) of each curve signifies the point of steepest downward slope in the release phase. The minimum of the second derivative (green) shows the ‘break’ point in each trace indicating the end of swelling and the onset of the release phase. Fits of the fast falling phase to the simplified Rayleigh-Gans equation are in black. Abbreviations: **ES** – extent of swelling; **tro** – time to release onset, **tsp** – time to the steepest point, **FP** – fraction of permeable osmolytes.

**Figure 5.**
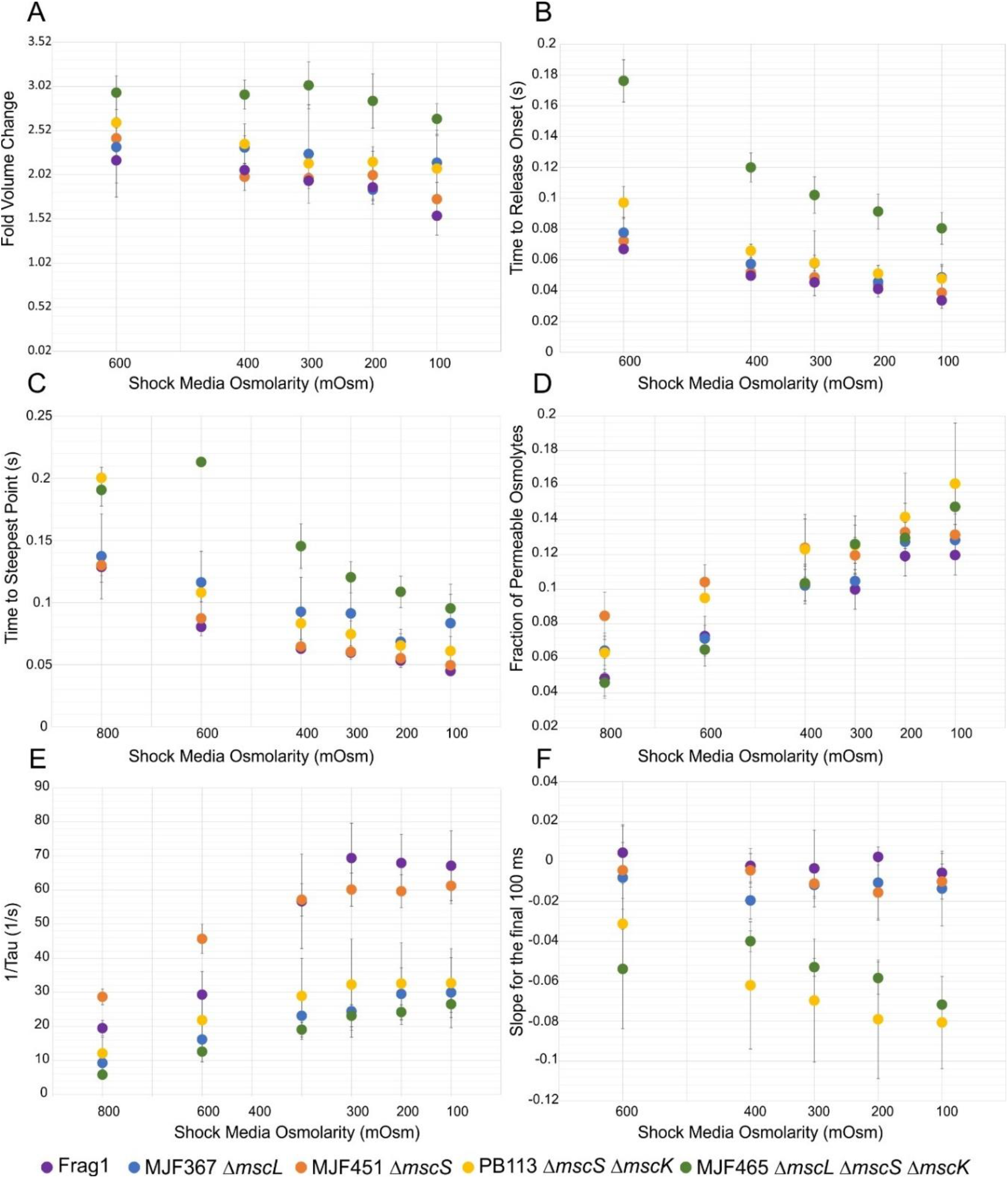
The parameters of cell swelling and osmolyte release kinetics determined from the stopped-flow traces. The explanation of parameters is given in Figure 4 and the color coding of strains is show at the bottom.

**Figure 6.**
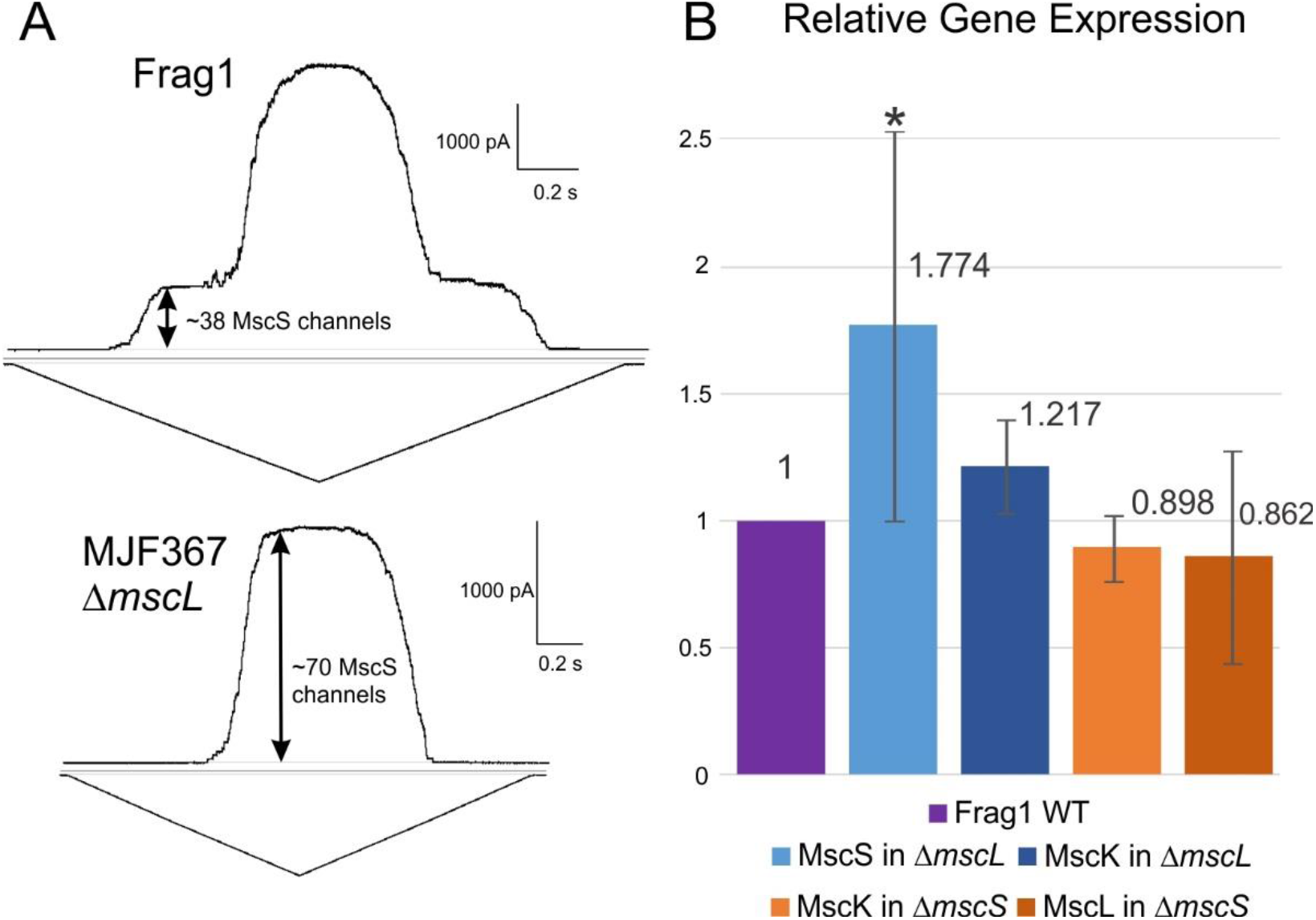
The size of the MscS population increases in the absence of MscL. (A) Patch-clamp population currents (top traces) recorded at +30 mV(pipette) in response to saturating (100-200 mm Hg) triangular pressure ramps (bottom parts). The amplitude of current estimates the number of channels per patch. The statistics of 6 patches taken from each strain gave 38 ± 13 MscS channels in Frag1 and 70 ± 34 in MJF367, with the ratio of 1.83. This increase was found to be significant using a one-tail two-sample t-test assuming unequal variances with a p-value of 0.04. (B) Levels of mRNA for major channels in the absence of their counterparts based on qRT-PCR experiments. Six independent experiments gave the ratio of 1.77 ± 0.76. Statistically signifigant (p=0.02) expression level when compared to the Frag1 strain is represented with an asterisk.

### Supplemental Material

The Supplemental Material contains the following sections: (S1) t-test p-values for all osmotic viability assays; (S2) derivation of the simplified form of the Rayleigh-Gans equation; (S3) determination of the refractive index increment for several solutes, immersive refractometry data on single *E. coli* cells and calibration curves calculated for the scattering intensity during swelling and release phases; (S4) table of channel numbers and midpoint pressure values from patch-clamp; (S5) RT qPCR primer sequences and primer efficiency graphs; (S6) functional comparison of MJF465 cells expressing MscS or MscL, survival and stopped-flow; (S7) kinetic parameters of extracted from stopped-flow traces of MJF465 cells expressing MscS and MscL; (S8) functional comparison of MJF429 and PB113 strains, survival, stopped-flow, and kinetic parameters; (S9) patch-clamp traces illustrating the degree of inactivation for WT MscS and two non-inactivating mutants, G113A and G168D; (S10) Western Blot and immunostaining micrographs illustrating expression levels of WT and G168D MscS; (S11) kinetic parameters of extracted from stopped-flow traces of MJF465 cells expressing G168D MscS; (S12) survival of MscS non-inactivating mutants compared to the relevant strains; (S13) patch-clamp traces comparing all strains and plasmids under the same pressure protocol illustrating ‘lingering’ channel activities at their activation thresholds for Frag1, MJF451, and MJF367 and slow closure of the non-inactivating G113A mutant, all relative to WT MscS.

## Results

### 1. Osmotic survival of the chromosomal deletion mutants

Since the expression level of mechanosensitive (MS) channels is important for the osmotic rescuing function, we chose to use chromosomal deletion mutants in order to single out activities of specific channels at their native expression density. Cells were grown in 1200 mOsm LB (supplemented with NaCl), shocked into media of lower osmolarity, diluted, and plated. The fractions that survived at different down-shocks are shown in Figure 1 in comparison to the no shock control (1200 mOsm).

When observing Frag1 with the full complement of channels in comparison to the triple knockout MJF465, the difference in survival becomes most evident for the down-shocks to 600 mOsm or lower. At 400 mOsm, MJF465 survival falls to 10%, while Frag 1 maintains 70-80% survival; the extreme down-shock to 200 or 100 mOsm kills most cells in both populations. Interestingly, PB113 (Δ*mscS* Δ*mscK* Δ*recA*) fares worse than the triple knockout at 800 and 600 mOsm (moderate shock) despite having MscL, the last resort emergency release valve. Furthermore, the MJF367 strain lacking MscL actually outperforms the parental strain Frag1 with all the channels present, as seen at 600 mOsm. Paradoxically, despite having the MscL gene that proved toxic in PB113 and lacking the MscS gene, the MJF451 (Δ*mscS*) strain still carrying *mscK* survives quite similarly to Frag1. This indicates that it is not the presence or absence of the channels alone that is the definitive reason for survival, but rather the functional interaction between these channels. Importantly, the MJF429 strain (Δ*mscS* Δ*mscK*), from which PB113 was derived by deletion of *recA*, survives similarly to Frag1. This shows that the absence of RecA creates a new context for MS channel function and interaction that impairs cell survivability under osmotic pressure. A similar context to PB113 for the function of MscL alone exists in MJF465, the triple KO mutant. Over-expressed WT MscL on plasmids does not rescue the population in osmotic viability experiments completely, whereas WT MscS under the same expression regime fully restores and even improves survival (see Section 4 and supplemental Figure S6). Reintroduction of WT MscS on a plasmid fully complements the ‘fragile’ survival phenotype of the PB113 strain as shown in supplemental Figure S8.

### 2. Stopped-flow experiments

We begin with a qualitative description of the light scattering traces obtained in rapid mixing experiments comparing Frag1 carrying the full complement of channels with MJF465 (Δ*mscS*, Δ*mscK*, Δ*mscL*) (Figure 2A). After a short mixing period, all traces show a stretch of nearly linear decrease of scattering followed by a steeper and larger drop that resolves in a plateau-like final level. According to the Raleigh-Gans equation (Supplement S2 and S3), the intensity of scattering is approximately proportional to the square of the ratio of refractive indexes inside and outside the cell. Because the refractive index inside the cell is directly proportional to the mass fraction of non-aqueous compounds (‘dry weight’) and is essentially independent of the composition (see (Koch, 1961; Koch et al., 1996) and supplemental Figure S3A), the scattering intensity is roughly proportional to the square of the total concentration of internal substances. Some of the small intracellular osmolytes are permeable through the channels, but most of the mid-size osmolytes and macromolecules are not. We attribute the decrease of scattering to two sequential processes: cell swelling and osmolyte release. *E. coli* cells remain turgid under 1200 mOsm growth conditions by pumping in osmolytes, yet the inner membrane always has some excess area (Buda et al., 2016), and it is expected that osmotic down-shock will additionally stretch the peptidoglycan and will allow for moderate swelling. The swelling dilutes the internal content and this leads to a gradual decrease of scattering upon mixing. Once the inner membrane completely unfolds, tension is generated that leads to channel activation. Therefore, following the period of initial swelling, the curve breaks and bends down with a steeper slope signifying the onset of osmolyte release. Comparison of essentially channel-free MJF465 with Frag1 shows that the swelling phase proceeds with similar kinetics, apparently because of similar water permeabilities of cell envelopes, but the release phase in the triple knockout is substantially delayed and the downward slope is shallower. The absence of major mechanosensitive channels indicates that the osmolyte release from MJF465 is mediated not by dedicated release valves, but rather by non-specific membrane breaks. The extended length of the swelling period in MJF465 and the magnitude of swelling reflected by the depth of the initial phase of the scattering trace are the signatures of irreversible cell damage directly correlating with less than 20% survival of the population beyond a 600 mOsm downshift (See Figure 1).

Moreover, the length and depth of the cell swelling phase are not the only indicators of death. Importantly, the final level of scattering for Frag1 stabilizes within 200 ms (a ‘hard stop’), indicating fast resealing of the membrane. In contrast, for both the triple knockout and PB113 (Figure 2B), the traces do not stabilize over the duration of the measurement. Under strong shocks, the Frag1 traces stabilize at the normalized scattering level of 0.63-0.65, whereas the traces of both MJF465 and PB113 approach 0.55-0.6, signifying a higher fraction of released internal osmolytes. The presence of MscL in PB113 is evidenced by the shortened swelling phase and steep drop of scattering, signifying fast release of permeable intracellular osmolytes. However, PB113 with MscL as the major release valve and lacking MscS and MscK does not survive mild shocks (see Figure 1), showing an even more toxic phenotype than the absence of all three channels in MJF465. However, at extreme shocks beyond 300 mOsm, MscL still slightly alleviates the lethal effect as compared to MJF465. The osmolyte release traces in Figure 2 show that MscL clearly curbs the swelling, but apparently fails to properly terminate the osmotic response. The absence of a ‘hard stop’ appears to be another signature of compromised viability.

We next consider the strains that behave comparably to or better than Frag1 in osmotic survival (Figure 1), MJF367 (Δ*mscL*) and MJF451 (Δ*mscS*), in Figure 3A and 3B respectively. The deletion of MscL slightly increases the time to steepest point compared to the full complement of channels in Frag1, which shows the fastest release. But paradoxically, MJF367 surpasses Frag1 in osmotic survival while MJF451, with MscL and another low-threshold channel MscK, behaves similarly to Frag1. Both strains show a relatively stable end-level, apparently conferred by low-threshold channels, that is comparable to the ‘hard stop’ in Frag1. In the case of MJF367, the low-threshold channels MscS and MscK are both present, whereas in the case of MJF451 only MscK is present. This observation points to MscK having a functional role of disarming MscL and terminating the permeability response even in the absence of MscS.

### 3. Quantitative treatment of scattering traces

According to the Rayleigh–Gans equation (Koch, 1961; Koch et al., 1996), the angle-dependent scattering of a cellular suspension depends mainly on the effective cell size and the ratio of refractive indexes inside and outside the cells. Supplemental Figure 3A shows the dependence of the refractive index on mass fraction for eight chemically different solutes. For all of them, the data points lay close to the common straight fitting line with a slope of 0.00015 (mg/ml)^-1^ and the intercept of 1.331, which corresponds to the refractive index of water. The fitting line, therefore, links the refractive index inside the cell with the combined mass fraction of all non-aqueous components (dry weight). Supplemental Figure S3B shows phase contrast micrographs of three Frag1 cells pre-grown in 1200 mOsm and suspended in their growth medium supplemented with 35, 42 and 50% of Ficoll 400. As seen from the intensity profiles, the brightness of the cell changes from low in 35% Ficoll to high in 50%, whereas in 42% Ficoll the cell blends into the background. The cell appearance in phase contrast depends on the same ratio of refractive indexes as the scattering (Ross, 1967). According to direct refractometer measurements, the high-osmotic LB supplemented with 42% Ficoll has a refractive index of 1.385, and so the cytoplasm must be the same under 1200 mOsm growth conditions. From the graph in Supplemental Figure S3A, this index corresponds to the combined weight fraction of intracellular constituents of about 350 mg/ml. Starting from this initial point and using the full form of the Rayleigh–Gans equation we have predicted the changes of scattering intensity for the separate processes of cell swelling and osmolyte release by varying either cell volume and assuming no osmolyte exchange (Supplemental Figure S3C) or internal osmolyte mass fraction (Supplemental Figure S3D) in cells while keeping the volume constant. These calibration curves predict that the drop of scattering during the swelling phase by 5%, for instance, corresponds to a 1.7-fold increase of cell volume. The subsequent drop of scattering to 65% of the initial value would signify a loss of about 18 % of internal osmolytes. For extracting the characteristic time of osmolyte release (τ), the simplified approximation of the Rayleigh–Gans equation was derived (Cetiner et al., 2017) and used here for fitting the rapidly falling release phase of the scattering traces.

Figure 4A compares the time courses of scattering recorded on each of the five strains in response to an equal 1200 to 300 mOsm downshift. It shows a spectrum of responses between Frag1 (WT) and MJF465 (triple knock-out strain). Figures 4B and C provide the details of the fastest protective response observed in Frag1 in comparison with the delayed response of unprotected MJF465 cells. They differ in the extent of swelling, the time to release onset, time to the steepest point as well as in the rate of release (1/tau) determined from the fit (black curve). The strains also differ in the scattering level at the end of the 0.3 s trace, which measures the fraction of permeable osmolytes, and in the slope at the end reflecting the residual permeability. The first derivative of each trace (blue line) shows the minimum that marks the point of the steepest slope, whereas the second derivative (green line) has its minimum at the breaking point signifying the onset of release.

The detailed comparison of extracted swelling/release characteristics is shown in Figure 5 as a function of the down-shock magnitude. MJF465 swells the most and takes the longest to release, followed by PB113, with Frag1 reaching the release phase the quickest and MJF367 and MJF451 very close to Frag1 (Figure 5B). This illustrates the action of low threshold channels versus high threshold or lack of channels. The extent of swelling (fold of volume increase before the onset of release, Figure 5A) varies from 1.5 for Frag1 to ~3 for MJF465. We should note that the estimated swelling of fully protected cells in the stopped-flow experiments is considerably larger than previously observed on single *E. coli* cells (1.1-1.2) (Buda et al., 2016). It could be caused by the slower rate of dilution in microfluidic devices, which was shown to be a critical shock parameter (Bialecka-Fornal et al., 2015), or by longer observation times and a possible overlap between the swelling and release phase, which would blunt the extent of swelling. The extent of swelling of unprotected MJF465 cells (~3) apparently reflects the expansion of cell volume beyond the elasticity limit, leading to irreversible damage of the cell wall and blebbing under excessive turgor pressure. It was shown previously (Reuter et al., 2014) that rupturing of the peptidoglycan occurs in the time scale of 100-200 ms, but cell lysis resulting in formation of protein-depleted ghosts or complete phase-contrast disappearance takes seconds. Based on the fraction of permeable osmolytes (Figure 5D), the trace illustrating lethal shock in Figure 4C does not imply complete bursting of cells in the 300 ms timescale.

The time to steepest point (Figure 5C) was taken from the point at the minimum of the first derivative, which indicates the point of the fastest release and maximum channel activity. Again, Frag1 is the fastest to reach this point and MJF465 takes the longest, with the exception of the 800 mOsm mild shock that is fit to the swelling because there was little to no osmolyte release. The trend for the rest of the strains from fastest to slowest is MJF451, PB113, and MJF367. This corresponds to the activation of the mixed population of MscL and MscK, MscL alone, and MscS alone, respectively. At 800 mOsm, PB113 takes the longest to reach maximal channel activity because the internal osmotic pressure has to reach the high threshold for MscL (12-14 mN/m). The fraction of permeable osmolytes (Figure 5D) is taken from the initial scattering intensity to the point where the traces level off. At high-osmolarity shocks MJF465 is the least permeable because it does not release through dedicated channels and mostly just swells, while the MJF451 and PB113 strains, containing large MscL and lacking MscS, are the most permeable. At stronger (lower osmolarity) shocks Frag1 fares the best with the lowest fraction of permeable osmolytes. As the ending normalized light scattering level indicates, MJF465 is more permeable because it lyses and PB113 is the most permeable because of the continued leakage and the lack of a hard-stop. 1/τ is the rate of release from the fit of the light scattering to the modified Rayleigh-Gans equation (Figure 5E). Frag1 and MJF451 have the fastest rate of release, followed by PB113, MJF367, and finally MJF465 with the slowest release. This directly corresponds to the presence and conductance of available channels. The slope for the final 100 ms (Figure 5D) signifies the downward ‘creep’ at the end of the trace. These are close to zero for Frag1, MJF451, and MJF367 and more negative overall for PB113 and MJF465. In conclusion, PB113, with MscL only, shows traces that clearly curb the swelling via osmolyte release, but lack the ‘hard stop’ indicative of membrane resealing. MJF367, with MscS only, reaches the release phase earlier and stabilizes after the initial release phase. MJF451 survives well apparently because of MscK. The presence of a low-threshold channel appears to be necessary for termination of the permeability response and fast resealing of the membrane.

### 4. The rescuing capacity of MscL in the absence of MscS and MscK depends on the cellular context

The observed difference in osmotic survival between *mscL*-carrying strains MJF429 (Δ*yggB*, Δ*mscK*) and PB113 (Δ*yggB*, Δ*mscK*, Δ*recA*) is drastic despite the fact that the strains are almost isogenic. The data presented above (Figures 1–3) show a direct correlation of a ‘good survivor’ phenotype with the presence of a stable end-level (‘hard stop’) on the stopped-flow traces for four strains. For this reason, we conducted an additional series of stopped-flow experiments to compare the MJF429 and PB113 strains side-by-side. The results are compiled in supplemental Figure S8. Panel A summarizes the osmotic survival of these strains in comparison with positive and negative controls. The stopped-flow traces (panel B) qualitatively illustrate that although both strains show very similar kinetics of swelling and release, they differ in their ability to properly reseal the membrane. At high shocks, PB113 releases more osmolytes with a downward ‘creep’ that is characteristic of incomplete membrane resealing while MJF429 shows an almost perfect ‘hard stop’ at the end, signifying a more complete resealing. Indeed, the time to the steepest point (panel C) and the rate of release (1/Tau, panel E) extracted from these traces are essentially indistinguishable. The data show a small difference in the fraction of permeable osmolytes (panel D) but noticeably different slopes at the end (panel F). PB113 clearly does not reseal well and continues to leak, likely through MscL channels that remain active (see section 6 below). The more stable end-level in MJF429 can be attributed to the activities of the remaining MscS paralogs YnaI, YbdG, YbiO, and YjeP, which may ‘deflate’ the cell at the end of the shock response and thus pacify ‘leaky’ MscL. The absence of RecA in PB113 may lead to this difference, perhaps due to expression levels. It is known that RecA is the main component of the cellular SOS stress response, which activates multiple genes involved in DNA repair. Additionally, RecA has been implicated in swarming motility (Gomez-Gomez et al., 2007), adhesion to surfaces (Costa et al., 2014) and biofilm formation (Kaushik et al., 2022), which may affect cell envelope mechanics and expression of mechanosensitive channels. We hypothesize that deletion of *recA* may silence the expression of residual MscS-like channels, which normally (in MJF429) would provide additional permeability and quickly deactivate MscL after the shock. In support of this requirement of additional permeability, we also show that re-expression of WT MscS in PB113 completely restores the ‘good survivor’ phenotype (Figure S8A). The question regarding the effects of RecA deletion on cell mechanics would require a special study.

Furthermore, we compared the rescuing ability of MscS and MscL re-expressed in the commonly used MJF465 strain. Supplemental Figure S6A presents the stopped-flow traces obtained in MJF465-MscS and MJF465-MscL strains after full induction with IPTG (1 mM, 30 min). It is clearly seen that cells expressing the high-threshold MscL experience a delayed release compared to those expressing MscS. Additionally, the slope over the final 100ms of the measurement was higher for MscL at stronger shocks.

Panel B shows that the osmotic survival of MJF465-MscL is compromised compared to Frag1 despite the higher plasmid expression levels. MscS alone imparts osmotic viability to the triple knock-out strain at a level similar to Frag1, whereas with MscL alone, the viability at moderate shocks is reproducibly 20-30% lower. At high shock magnitudes, overexpressed channels do impart higher viability than Frag1. The 70-80% survival with MscL is consistent with the original results reported by Levina (Levina et al., 1999) and more recent data by Chure (Chure et al., 2018). Supplemental Figure S7 depicts the kinetic parameters of osmolyte release from five strains extracted from stopped-flow traces. The noticeable difference is a higher negative slope of MJF465-MscL traces (panel F) that correlates with lower osmotic survival.

### 5. Expression of MscS is MscL-dependent

The remaining question is: why does MJF367 (Δ*mscL*) outperform the parental strain Frag1 in terms of viability? Fast osmolyte release from the Δ*mscL* strain and the high survival rate in osmotic experiments prompted us to look at the density of channels in standard patches (r =1.4 μm) and at the expression level of the channels at the mRNA level using qRT-PCR. Both methods gave a consistent 1.8-fold increase of the MscS population indicating that MscS is over-expressed in the absence of MscL (Figure 6).

We also checked the expression levels for MscK in both MJF367 and MJF451 and the expression of MscL in MJF451, but saw no significant change in expression. The data indicate a compensatory increase of MscS expression when MscL is deleted. The level of MscK mRNA also tends to increase, but not as dramatically. It was previously reported that MscS and MscL are constitutively expressed under all growth condition but up-regulated 2 to 4-fold when the culture enters the stationary phase and especially when grown in high-osmotic media. Their expression was suggested to be controlled by the σ^S^ transcription factor (RpoS) that activates many stress genes (Stokes et al., 2003). The RegulonDB database lists 39 σ^S^ -dependent operons, four of which are osmotically-regulated (*osmC, E, Y and otsBA*) and 38 σ^S^ -dependent genes, but *mscS* is not among them. Instead, the RegulonDB database indicates *mscS* is regulated by an operon named mscSp, which is predicted by the *E. coli* promoter prediction tool iPro70-FMWin to be a σ^70^ promoter sequence (Rahman et al., 2019).

While the exact regulation for these genes is still unclear, our results show that the expression machinery of *mscS* (and possibly *mscK*) in some way ‘feels’ the absence of MscL. This suggests a possible cross-talk between the low-threshold channel expression, especially MscS, and cell mechanics. We cannot exclude that *mscL* mRNA has an effect on *mscS* transcription, but more likely it is the absence of functional MscL channel that produces some stress that is compensated by a higher level of MscS. Another possibility is that in the absence of MscL the extremely crowded inner membrane has more vacancies to accommodate the extra amount of MscS, and for this reason the continuously translated *mscS* mRNA has a longer life and a higher copy number.

### 6. Patch-clamp dissection of the closing process

Patch-clamp protocols mimicking the pressure changes that cells experience under osmotic down-shock were performed to further examine functional interactions between the channels and to look specifically at the behavior associated with channel activity termination. For a simple channel that opens and closes in response to tension, the presumption is that when the tension in the bacterial membrane drops down to the threshold for that specific channel, then that channel population closes and can no longer meaningfully contribute to lowering the membrane tension. Thus, if this is the only channel population present in the membrane, the remaining tension cannot be dissipated any further and instead remains fixed near the threshold for the active channel. As a specific example, if MscS, MscK and other low-threshold channels are absent from the membrane, then the osmotic shock would activate MscL and, upon the end of release phase, the tension would stabilize at ~10 mN/m, just below MscL’s threshold. The question then becomes: how do high- and low-threshold channels behave at their own tension thresholds and, if they are not redundant, what is their unique contribution to the permeability response?

We see that MscL and MscS display markedly different closing behaviors at their threshold tensions. As illustrated in Figure 7A, once the full population of channels in Frag1 is opened and the pressure is slowly ramped down to 0 mmHg (red), two distinctive current waves are observed.

**Figure 7.**
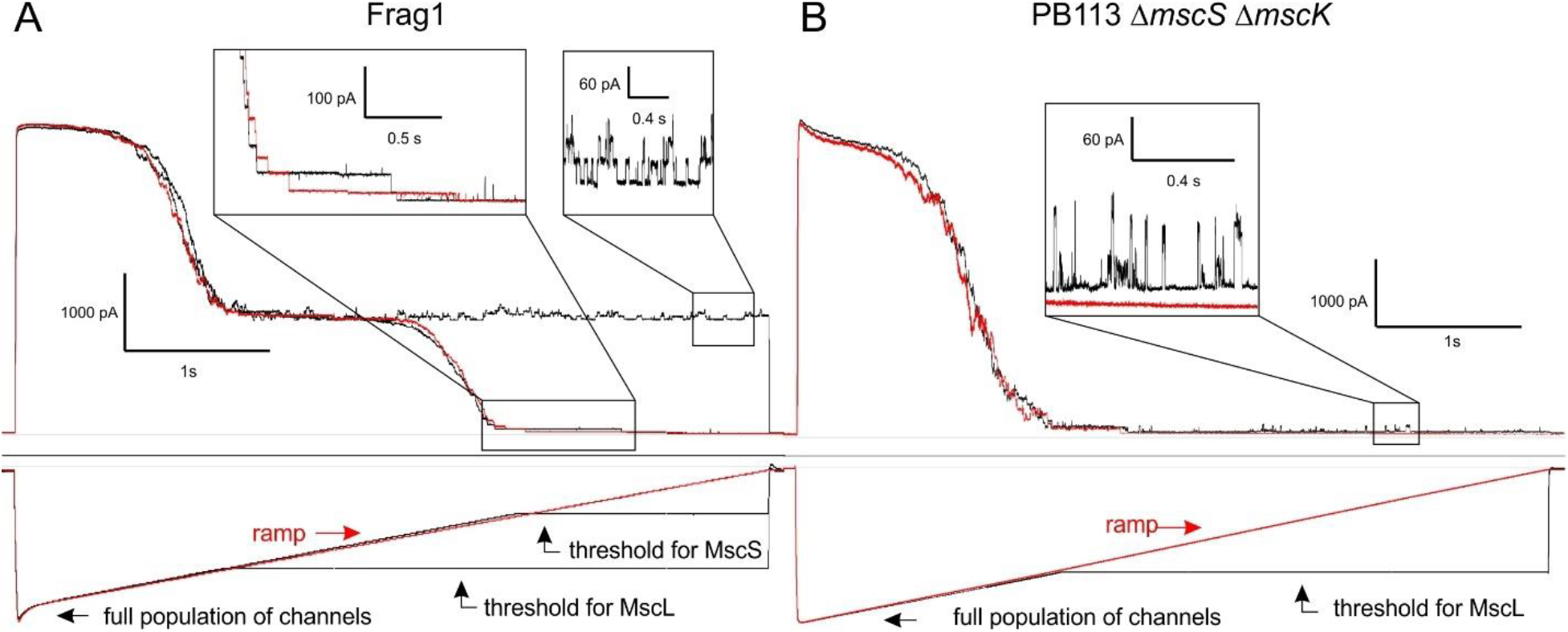
Traces of Frag1 (A) recorded under a fast activation step and a slow deactivation regime to observe the behavior of each population of channels. When the ramp halts at the activation threshold for MscL, the visible fraction of the MscL population stays active, producing constant flickering channel activity. When the ramp halts at the threshold for MscS, the activity of the small-conductance channel population continues to terminate and completely inactivates. Traces of PB113 (B) under the same protocol show residual MscL channel activity in the absence of a low-threshold channel. All recordings are done at +30 mV pipette potential.

The first wave reflects the closing MscLs, followed by the second wave of closing MscSs. If the ramp is held at the end point for the MscL population (black), i.e., the threshold for the MscL channel, then there is visible flickering of MscL as it opens and closes on top of the fully open MscS population (right box). This residual MscL activity is especially evident in the PB113 (Figure 7B). In contrast, when the ramp is stopped at the MscS threshold, the MscS channels continue to close and the patch becomes completely silent through MscS inactivation, despite the continued application of pressure.

### 7. Low threshold or inactivation, what is more important?

The differing closing phenotypes between MscL and MscS is due largely to MscS’s ability to adapt and close under constant tension and to inactivate upon closing (Cetiner et al., 2018; Kamaraju et al., 2011). Inactivation thus appears to be a key element of proper termination of the osmolyte release sequence. To test this hypothesis and probe the impact of inactivation on osmotic viability, we performed a set of experiments in which we compared the MJF465 strain expressing G168D, the non-inactivating MscS mutant (Koprowski et al., 2011; Rowe et al., 2014), with the same strain expressing WT MscS. Patch-clamp data comparing inactivation properties of non-inactivating MscS mutants with WT MscS obtained using the pulse-step-pulse protocol (Akitake et al., 2005) can be found in Supplemental Figure S9. Both channels, WT MscS and G168D MscS are 6-His tagged on their C-termini and their expression and location in bacterial cells can be reliably determined by in-situ immunostaining (Supplemental Figure S10B). Based on immunostaining and Western Blot (Supplemental Figure S10A), we determined that under standard induction conditions (1 mM IPTG, 30 min), both WT and G168D MscS are comparably expressed. The plasmid containing the G168D MscS was transformed into PB113 in order to compare the midpoint of the mutant MscS to the midpoint tension of the native MscL which is taken as 12 mN/m. The midpoint ratio for G168D is 0.54 +/- 0.01 and the tension is 6.4 +/- 0.1 mN/m, which is similar to the literature values for WT MscS at 0.6 for the midpoint ratio and 7.2 mN/m for the tension (Belyy et al., 2010). From the character of release traces (Figure 8A), one can see that the non-inactivating mutant opens at the same degree of swelling and in the beginning of the release phase it almost re-traces WT MscS. However, then G168D traces deviate from WT MscS and do not exhibit a clear hard stop. The fitted parameters can be found in Supplemental Figure S11. The viability data indicate that G168D MscS does not rescue cells as well as WT MscS (Figure 8B). Osmotic viability for another non-inactivating MscS mutant G113A (Akitake et al., 2007), also inferior in terms of osmotic recuing, can be found in Supplemental Figure S12. We conclude that the ability to inactivate is critical for proper termination of the massive permeability response.

**Figure 8.**
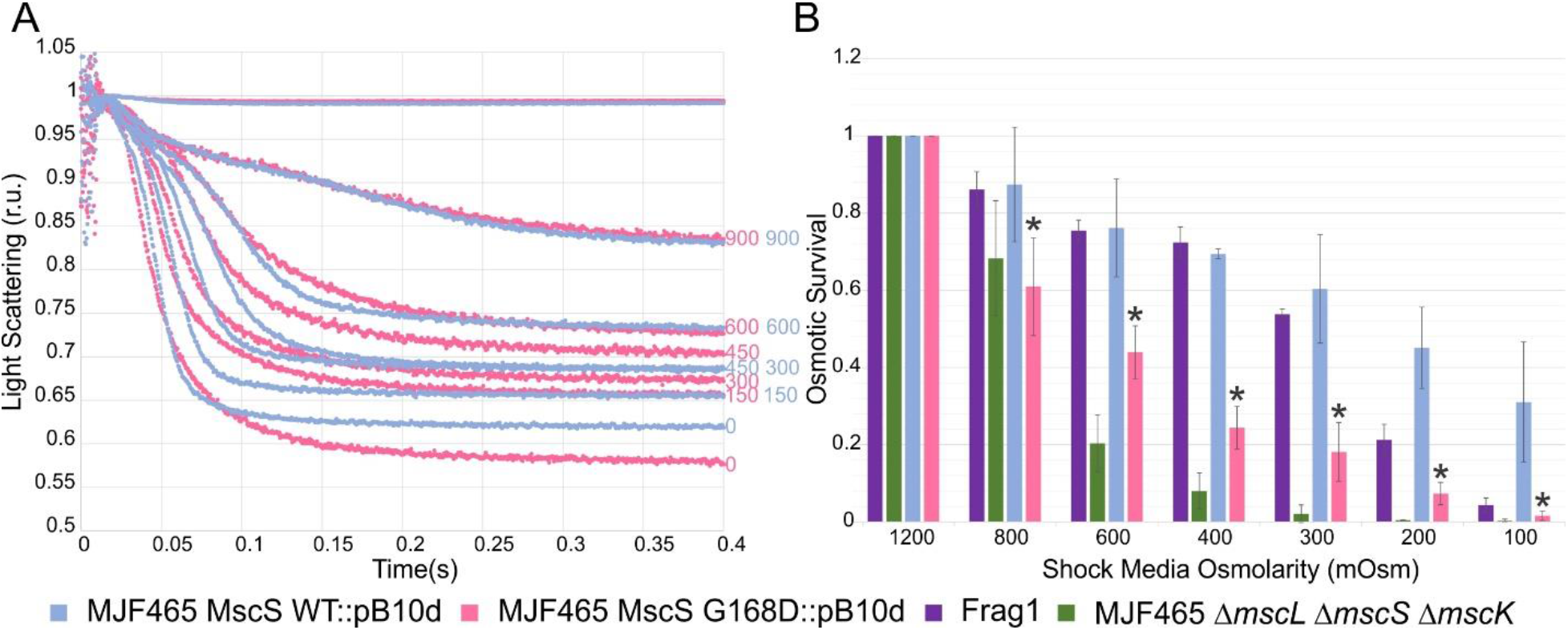
Comparison of MJF465 expressing WT MscS and non-inactivating G168D MscS on plasmids. Stopped-flow traces (A) and a bar graph of osmotic survival shown together with Frag1 and empty MJF465 (B). Statistically signifigant (p<0.05) survival values of G168D MscS compared to WT MscS are represented with an asterisk. A full table of all p-values can be found in Supplemental Table S1.

Additionally, patch-clamp traces reveal an extremely slow closing rate for G168D MscS when pipette pressure is dropped to its activation threshold (Figure 9). To achieve closing of the population within 4 s, the pressure had to be 30 mm Hg below the activation threshold. Patch-clamp traces of a similar protocol to Figure 9 can be found in Supplemental Figure S13 for Frag1, PB113, MJF451, MJF367, MscS WT on plasmids, and G113A.

**Figure 9.**
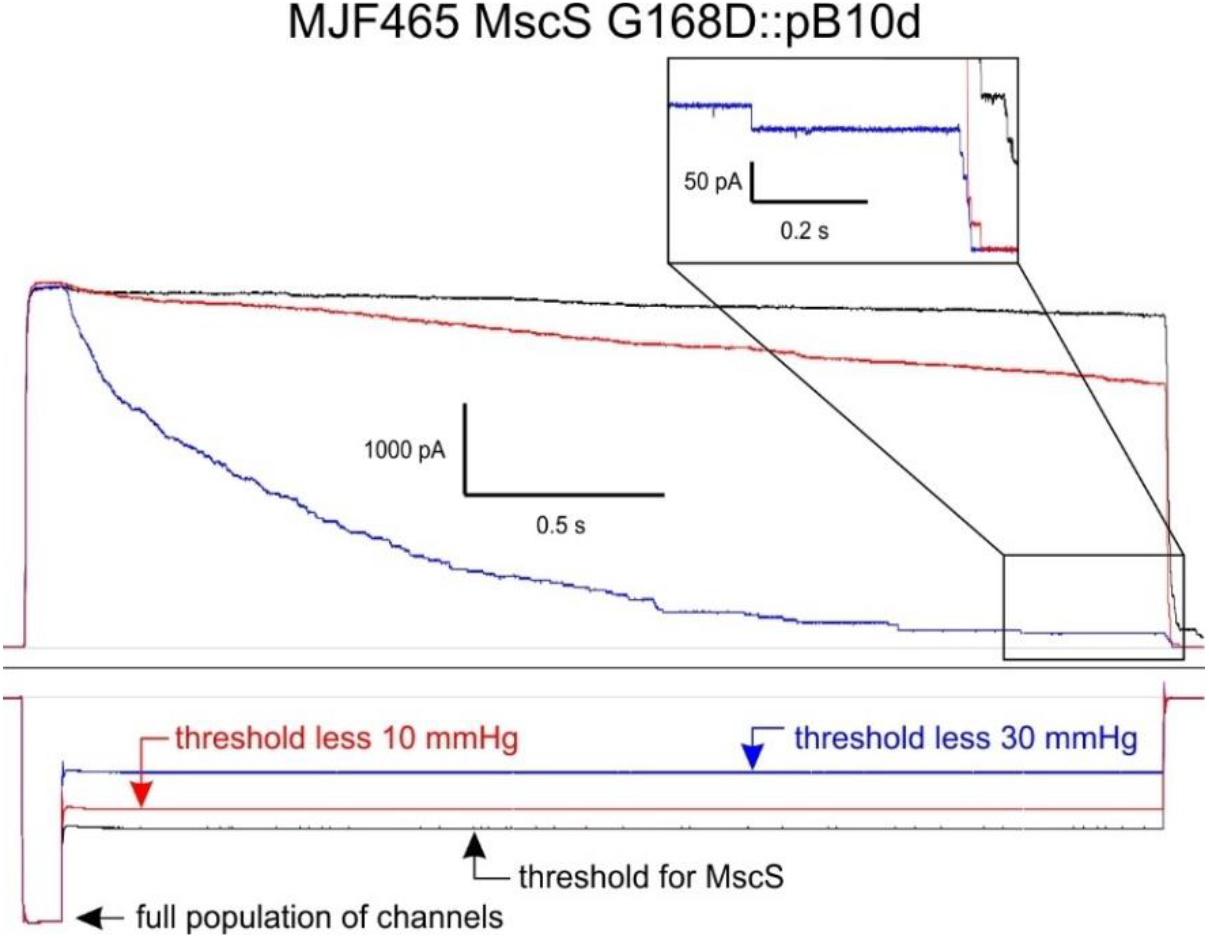
Patch-clamp traces from G168D MscS recorded at +30 mV in the pipette. The lingering of MscS channels is clearly visible even as the pressure is dropped to 30 mmHg below the opening threshold for MscS.

Importantly, besides delayed closing, the inactivation process of G168D is also resistant to increased crowding (Rowe et al., 2014), which apparently produces lingering activity. Slow closing and impaired inactivation seemingly make G168D MscS unfit for the role of permeability terminator device.

## Discussion

The very first observation that *E. coli* instantly expels its entire pool of free amino acids under osmotic down-shock was reported in 1962 (Britten and McClure, 1962). The electrophysiological discovery of mechanosensitive channels in the *E. coli* envelope pinpointed the nature of osmolyte release sites (Martinac et al., 1987). Early patch-clamp surveys of the *E. coli* cytoplasmic membrane and proteo-liposomes reconstituted with inner membrane protein fractions (Berrier et al., 1996; Sukharev et al., 1993) described three major phenotypical classes of MS channels differing in conductance and activating pressure: large (MscL, 3 nS), small (MscS, 1 nS) and mini (MscM, 100-300 pS). This variety was interpreted as the capacity of the system to respond to different shocks in a graded manner, activating mini- and small channels with the lowest thresholds first, and then, as necessary, engaging the large channel at near-lytic tensions. At that time, it was unclear how fast the release should be under extreme shocks and what the fold permeability increase should be in order to rescue cells. The data presented above provide some of these numbers. Upon a 1200 → 200 mOsm downshift, the expressed population of MscS (Figure 8), once activated, releases 90% of permeable osmolytes within ~25 ms, rescuing about half of the cell population. A simple estimation based on the energy of MscS opening transition (ΔG_C→O_ ≈ 24 kT) (Akitake et al., 2005; Cetiner et al., 2018) shows that at tensions that activate the majority of the population, the open probability, and respectively the permeability, increases by at least 9 orders of magnitude relative to the resting state. In the present paper we address the question of how the activities of large- and small-conductance channels rescue the cells and more specifically how the channels cooperate in terminating this immense permeability response.

The main findings show that MscS and MscL, the two major components of the osmolyte release system, are neither redundant nor equivalent in their function. The extent of cell swelling upon abrupt dilution appears to be the critical parameter that defines whether the cell will be irreversibly damaged, although this does not appear to be the only factor impacting survival. Both MscS and MscL are similarly efficient in reducing the swelling about two- to three-fold compared to unprotected cells. However, MscL, the large non-inactivating emergency valve, must be co-expressed with a low-threshold inactivating channel, otherwise it becomes toxic. We must emphasize that the conclusion about the toxicity of ‘lonely’ MscL and its inability to rescue cells was obtained in a specific setting, in the PB113 (Δ*mscS* Δ*mscK* Δ*recA*) strain, which revealed a leaky state of the cell after the shock. Similarly, expression of MscL in MJF465 (Δ*mscL* Δ*mscS* Δ*mscK*) does not rescue the population completely. Complementation of PB113 and MJF465 with WT MscS completely rescued the osmotically fragile phenotype. We did not observe, however, the leaky state in MJF429 (Δ*mscS* Δ*mscK*) suggesting that this double mutant may not provide the ‘clean’ background to single-out the lingering MscL activity after the shock. It is now clear that the role of early-activated MscS and MscK after the shock is to reduce tension in the membrane below the threshold for MscL in order to completely silence it. The ability to inactivate appears to be crucial for the low-threshold channel since a non-inactivating mutant of MscS does not osmotically rescue cells as effectively as WT MscS does. Inactivation of the lowest-threshold channel appears to be the last step in the termination of the osmotic permeability response.

We chose to conduct our experiments on a set of chromosomal deletion mutants (Levina et al., 1999), with the expectation that the remaining channels would be expressed at their native levels. This was the case with MscL and MscK as revealed by qRT-PCR; however, deletion of *mscL* resulted in a 1.8-fold increase of *mscS* mRNA, which correlated with the same increase of channel density in standard patches. This result strongly suggests a cross-talk between the osmotic/mechanical status of the cell and *mscS* transcription. This warrants a closer look at the *mscS* promoter, associated sigma factors attenuation/termination elements, and the biophysical mechanism of transcription activation. We found that MscS by itself is extremely effective as a rescuing osmolyte valve, and its overexpression results in osmotic viability that surpasses WT bacteria (Frag1). Higher than Frag1 viability was also observed when WT MscS is over-expressed from a plasmid (Figure 8B).

Early light-scattering (Koch, 1961; Koch et al., 1996) and refractometry (Valkenburg and Woldringh, 1984) measurements on bacterial suspensions laid the necessary foundation for our studies of osmolyte release. Our estimates obtained using immersive refractometry suggested that *E. coli* cells grown in rich 1200 mOsm media contain 350-400 mg/ml of non-aqueous solutes inside. This is consistent with the data reported by Cayley that the ratio of intracellular water volume to dry weight of a hypertonically-adapted cell is about 1.7 μL per mg of dry weight (Cayley and Record, 2004). This translates into approximately 400 mg/ml of total intracellular solutes. Our analysis of scattering traces shows that up to 18% of these non-aqueous contents are dispensable and small enough to permeate through mechanosensitive channels (Wood et al., 2001), thus relieving the mechanical stress, with the remaining larger molecules and polymers having a smaller contribution to the osmotic pressure.

The stopped-flow results shown in Figures 2–5 present a physical picture of what the kinetics of swelling and osmolyte release look like in the event of osmotic damage to unprotected cells as opposed to channel-mediated rescue. The interpretation of traces is based on the simple relationship where the scattering level is proportional to the square of the osmolyte concentration ratio inside and outside the cell. The first event seen on the scattering traces is elastic swelling of the cell until the point when sufficient tension is generated in the inner membrane to open the mechanosensitive channels and initiate osmolyte release. In the absence of the three major channels (MJF465 Δ*mscS*, Δ*mscK*, Δ*mscL*), the swelling phase is about two times longer than in Frag1. A greater extent of swelling is the most probable cause of irreversible damage. When the channels are present, they shorten the swelling phase substantially and the cell quickly switches to the release phase. Under extreme shocks the cells release 15-18% of their non-aqueous content in the form of small permeable osmolytes within ~20-30 ms. The fast release rate must outcompete the rate of water influx to curb excessive swelling and prevent lysis. Upon completion of the release phase, the permeability sharply decreases, producing a stable scattering level at the end of the trace (the hard stop) signifying a non-leaky state of the membrane. The hard stop correlates with the presence of low-threshold channels that are able to inactivate. In patch-clamp, inactivation of MscS gradually proceeds under moderate (near-threshold) tensions but is sharply augmented by the presence of crowding agents (Rowe et al., 2014). Thus, the reduction of cell volume that increases cytoplasmic crowding appears to be the main factor driving MscS inactivation and membrane resealing. The experiments with MJF451 (Δ*mscS*) strain indicated that MscK can be a functional substitute for MscS, which raises the question about its ability to inactivate. Previous reports (Li et al., 2002) suggested that MscK has even lower activation threshold than MscS, but there has been no mention of inactivation. Therefore, our result warrants further investigation of MscK inactivating behavior. Similarly, the conductive and gating properties of MscS paralogs ybdG, ybiO, yjeP and ynaI (Edwards et al., 2012; Schumann et al., 2010) need to be better understood.

In summary, in this work we have determined the contributions of MscL and low-threshold channels MscS and MscK in the mechanism of osmotic rescuing in bacteria. The stopped flow experiments provided clear correlations between the kinetics of osmolyte release and cell survival. Emphasis was placed on the mechanism of membrane resealing after a massive osmotic permeability response. We elaborated on the simple notion that a mechanosensitive channel cannot regulate membrane tension below its activation threshold. MscL, which provides the fastest release rate, ceases to function as a tension-regulating valve below 10 mN/m, and instead flickers and remains leaky at this tension. For this reason, a set of channels with different activation thresholds becomes beneficial for the gradual reduction of tension and intracellular pressure, respectively. Channels with lower tension thresholds pacify channels with higher thresholds. MscS and MscK are the two low-threshold valves that silence MscL and reseal the cell membrane. Inactivation of MscS appears to be the critical feature for the proper termination of the osmotic permeability response.

## Acknowledgement

This work was supported by NIH R01AI135015 to SS and by the National Science Foundation Graduate Research Fellowship Program under Grant No. DGE 1840340 to EM. Any opinions, findings, and conclusions or recommendations expressed in this material are those of the author(s) and do not necessarily reflect the views of the National Science Foundation. We would like to thank Dr. Samantha Miller (University of Aberdeen, UK) for providing most of the strains, Madhabi Majumdar for immersive microscopy data and Amy Beaven for expert assistance with qPCR.

