## Supplemental Material for "Mechanosensitive channel MscS is critical for termination of the bacterial hypoosmotic permeability response"

(S1) **p-values for t-tested osmotic viability assays**

| Comparison | 800 | 600 | 400 | 300 | 200 | 100 |
|---|---|---|---|---|---|---|
| MJF367 to Frag1 | 0.556 | ***0.006*** | 0.504 | 0.436 | 0.106 | 0.263 |
| PB113 to Frag1 | ***0.002*** | ***0.008*** | ***0.001*** | ***0.006*** | ***0.002*** | 0.160 |
| MJF465 to Frag1 | 0.088 | ***1.86E-9*** | ***6.52E-6*** | ***4.64E-9*** | ***0.002*** | 0.063 |
| WT MscS in MJF465 to Frag1 | 0.817 | 0.864 | 0.335 | ***0.044*** | ***2.16E-4*** | ***1.74E-4*** |
| WT MscS in PB113 to Frag1 | 0.981 | 0.528 | ***0.015*** | ***0.008*** | ***1.97E-5*** | 0.083 |
| WT MscL in MJF465 to Frag1 | ***0.008*** | ***9.12E-5*** | ***0.004*** | ***4.4E-4*** | 0.176 | ***8.62E-4*** |
| G168D to WT MscS | ***0.002*** | ***7.50E-7*** | ***1.23E-6*** | ***4.2E-13*** | ***5.12E-6*** | ***9.56E-5*** |
| G113A to WT MscS | 0.429 | 0.659 | 0.315 | ***0.006*** | ***0.034*** | ***8.88E-5*** |

Table S1: P-values from two-tail two-sample t-tests assuming unequal variances used to determine statistical significance between osmotic viability experiments. The strains compared are listed in column 1, all strains were considered for significance compared to Frag1 as were WT MscS and WT MscL. MscS mutants G168D and G113A were considered in comparison to WT MscS expressed on plasmids. Significant values are below 0.05 and are italicized and bolded. They are shown on bar graphs with asterisks.

(S2) **Equations for treatment of light scattering traces**

The stopped-flow technique combined with small-angle light scattering allowed us to observe the time course of bacterial osmotic responses to rapid dilution of the medium and quantify the swelling and release. The intensity of the scattered light depends on multiple parameters defined by the experimental setup. These are the angle of observation, the shape of bacterial cells, and the cytoplasmic composition. For a bacterial suspension, the light scattering intensity has been described by the Rayleigh-Gans approximation (Koch, 1961; Koch et al., 1996) where the main variables are the volume of cells and the ratio of refractive indexes for the cytoplasm and the surrounding media:

$$I = \frac{8\pi^4 r^6 \eta_0^4}{R^2 \lambda^4} \cdot \frac{\left[\left(\frac{\eta}{\eta_0}\right)^2 - 1\right]^2}{\left[\left(\frac{\eta}{\eta_0}\right)^2 + 2\right]^2} \cdot \nu I_0 V[1 + cos^2(\theta)] \cdot P(\theta) \qquad (1)$$

where $I$ is the intensity of the scattered light for a given direction and distance; $r$, the radius of the spherical equivalent of the actual particle (in our case, a rod-shaped bacterium); $\eta_0$, the index of refraction of the suspending medium; $\eta$, the index of refraction of the particle; $V$, the volume illuminated; $I_0$ the intensity of the incident light: $\nu$, the concentration of particles; $\theta$ the angle of observation; $R$ the distance of the detector from the sample; and $\lambda$, the wavelength of light in vacuum. The factor $[1 + cos^2(\theta)]$, depends on the angle of observation, $\theta$, relative to the forward direction of the illuminating beam. For a given physical setup, most of these factors can be combined into an empirical constant.

As was shown previously (Cetiner et al., 2017), with several empirical constants introduced, equation (1) can be transformed into this form:

$$I_r = I_b + Ar^6 \cdot \frac{\left[\left(\frac{\eta_w + \frac{3(S_p m_p + S_i m_i)}{4\pi r^3}}{\eta_0}\right)^2 - 1\right]^2}{\left[\left(\frac{\eta_w + \frac{3(S_p m_p + S_i m_i)}{4\pi r^3}}{\eta_0}\right)^2 + 2\right]^2} \qquad (2)$$

Here $I_r$ is the intensity of the recorded scattered light, $I_b$ is the background light component, $r$ is the radius of the spherical equivalent of the bacterium, $\eta_w$ is the refraction index of water, $\eta_0$is the refraction index of extracellular medium, $m_p$ and $m_i$ are the masses of the permeable and impermeable solutes respectively, $S_p$ and $S_i$ are the scaling coefficients between their concentrations and contributions to the refraction index of the medium ($S_p = d\eta/dC_p$ , $S_i = d\eta/dC_i$, equation 4) and $A$ is the scaling coefficient that combines parameters of instrumental amplification, system geometry and the angle of collected light.

The analysis of equation (2) shows that because the numerator is much smaller than the denominator, the scaling of $I_r$ is quadratic with respect to the term $\left(S_p m_p + S_i m_i\right)$ , and in the course of osmolyte efflux the refractive index difference between the cell and its environment is the major parameter and has a strong dependence on time. Under the assumption that contributions of permeable ($S_p$) and impermeable ($S_i$)osmolytes to the refractive index are the same (see Figure S1) and the cell volume changes are small, the time course of the scattering signal can be presented in the following form:

$$I = I_0 + S(m_i + m_p(t))^2, \qquad (3)$$

where $I$ is light intensity, $I_0$ is background light intensity, $S$ is scaling coefficient, $m_i$ is the mass of impermeable osmolytes, and $m_p(t)$ is the mass of permeable osmolytes as a function of time $t$. In the assumption of exponential release of the permeable fraction of osmolytes, equation 3 can be presented in this form

$$I = I_0 + S(m_i + m_{p0}e^{\frac{t_0 - t}{\tau}})^2 \qquad (4)$$

where $m_{p0}$ is the initial mass of the permeable osmolytes before the release, and $t_0$ is the time at which the exponential release has started.

(S3) **Calibration of light-scattering measurements**

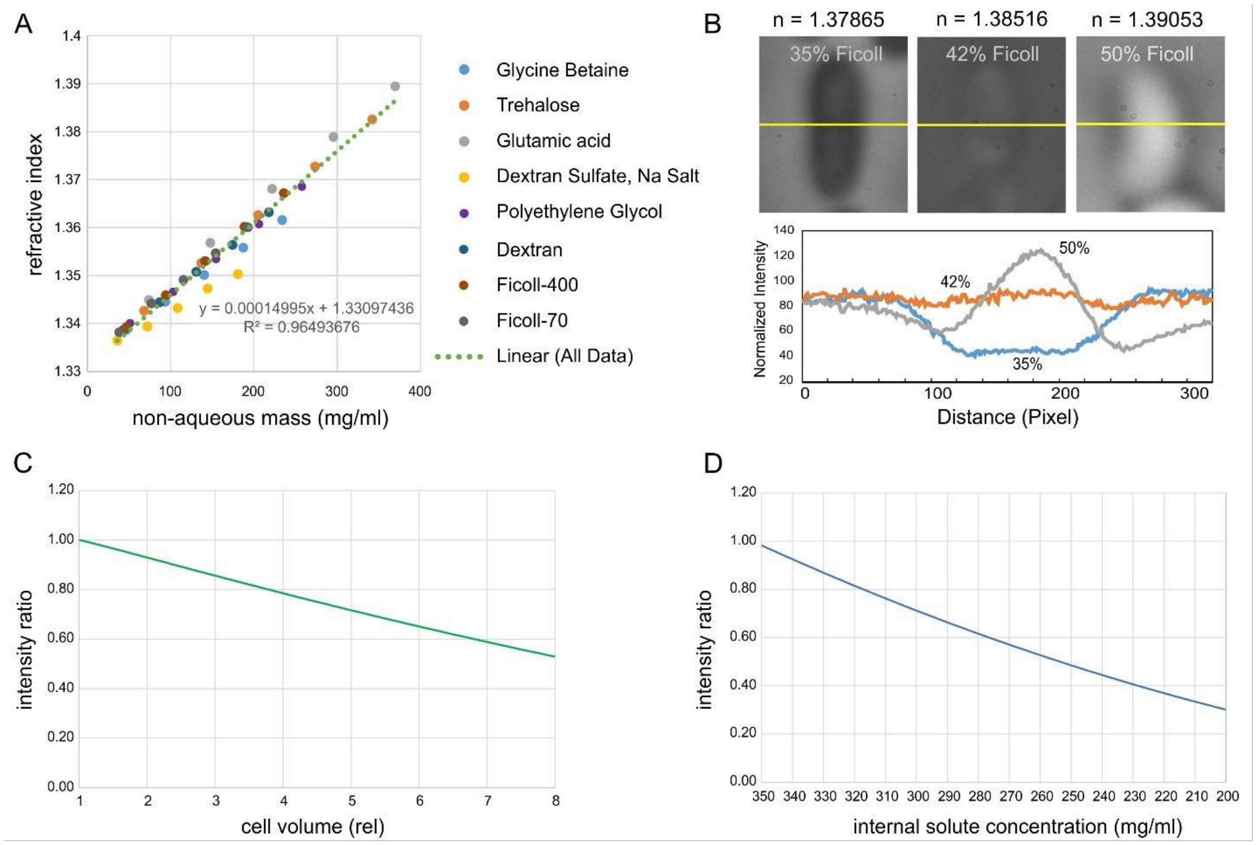


Supplemental Figure S3. Calibration graphs illustrating the dependence of the refractive index on concentrations of several solutes of varied chemistry (A). The linear fit for all substances gave us the common refraction increment of 0.00015. Previously published specific refraction increments vary from 0.000160 for NaCl to 0.000186 for human serum albumin (Barer, 1957). (B) The appearance of 1200 mOsm-adapted E. coli cells in phase contrast when submerged in high-density Ficoll 400 solutions of 35%, 42% and 50% by weight. The lower panel shows intensity slices along yellow lines for each case. The images show that at 42% of Ficoll the cell melds into the background indicating that the refractive indexes inside and outside the cell are equal. This principle of immersive refractometry has been previously described in (Barer, 1956; Liu et al., 2014; Valkenburg and Woldringh, 1984). (C) The scaling of scattering intensity calculated according to the full form of Rayleigh-Gans equation (equation 2) as a function of cell volume during swelling. (D) The scaling of scattering intensity as a function of internal solute concentration calculated for a cell in standard LB medium.

(S4) **Number of channels and gating pressures (in mm Hg) from patch-clamp measurements**

| Strain | Frag1 | MJF367 | MJF451 | PB113 |
|---|---|---|---|---|
| Number of MscS | 38 ±13 | 70 ±34 | | |
| Number of MscL | 69 ±18 | | 57 ±5 | 47 ±17 |
| MscS threshold pressure (mm Hg) | -78 ±15 | -62 ±13 | | |
| MscL threshold pressure (mm Hg) | -142 ±27 | | -121 ±35 | -118 ±14 |
| MscS midpoint pressure (mm Hg) | -89 ±17 | -76 ±12 | | |
| MscL midpoint pressure (mm Hg) | -163 ±28 | | -132 ±27 | -145 ±15 |
| MscS saturation pressure (mm Hg) | -107 ±17 | -91 ±15 | | |
| MscL saturation pressure (mm Hg) | -195 ±26 | | -165 ±29 | -199 ±28 |

Table S4: Full table of patch-clamp channel numbers and gating pressures. The estimated patch radii under the midpoint pressure were 1.4±0.1 µm and, correspondingly, patch areas varied between 11 and 15 $\mu m^2$.

(S5) **mRNA quantification**

RT-qPCR Primers (5’ to 3’)

MscK F: CCTGCTGCTGATTGCCTTCC

MscK R: AGCCTGTAGCAGTCAGCACC

MscS F: TACGCTCCAGCACATCCCAG

MscS R: CGCCTGAACGAACTTGGTGC

rpmA F: GCAAACAGAGTGTGGTCACGAC

rpmA R: CGTTCTGGCGGGTAGCATCATC

MscL F: ATTTGGCGGTGGGTGTCATTATC

MscL R: AGAGACAATCTTCCCGAATGCCG

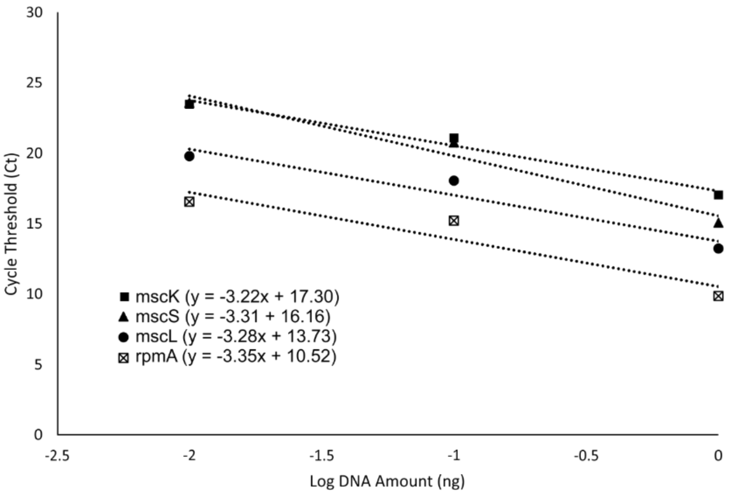


Supplemental Figure S5. Primer sequences and primer efficiency plots for RT qPCR experiments. qPCR analysis was conducted according to Livak (Livak and Schmittgen, 2001).

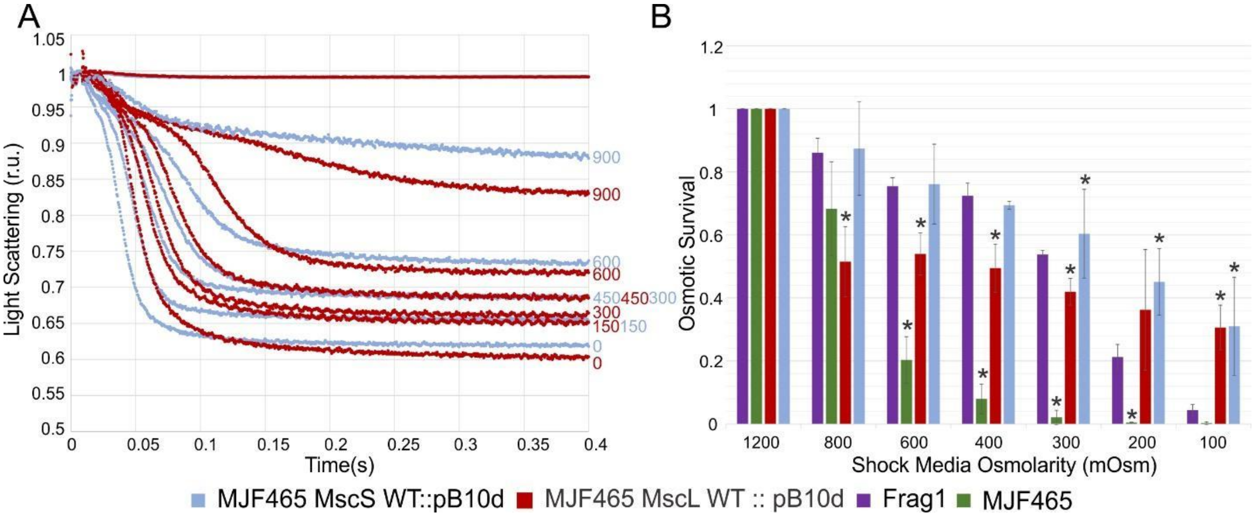


Supplemental Figure S6. Osmotic shock responses of MJF465 cells expressing either MscS or MscL.

(A) Stopped-flow traces comparing the kinetics of osmolyte release by individual populations of channels. (B) Osmotic survival of MJF465-MscS and MJF465-MscL strains in comparison with the positive (Frag1), negative (MJF465 empty) controls. The data show that MscS alone rescues the triple knock-out strain better than MscL alone. Statistically significant ($p<0.05$) differences in survival values when compared to the Frag1 strain are represented with an asterisk. The full table of all p-values can be found in Supplemental Table S1.

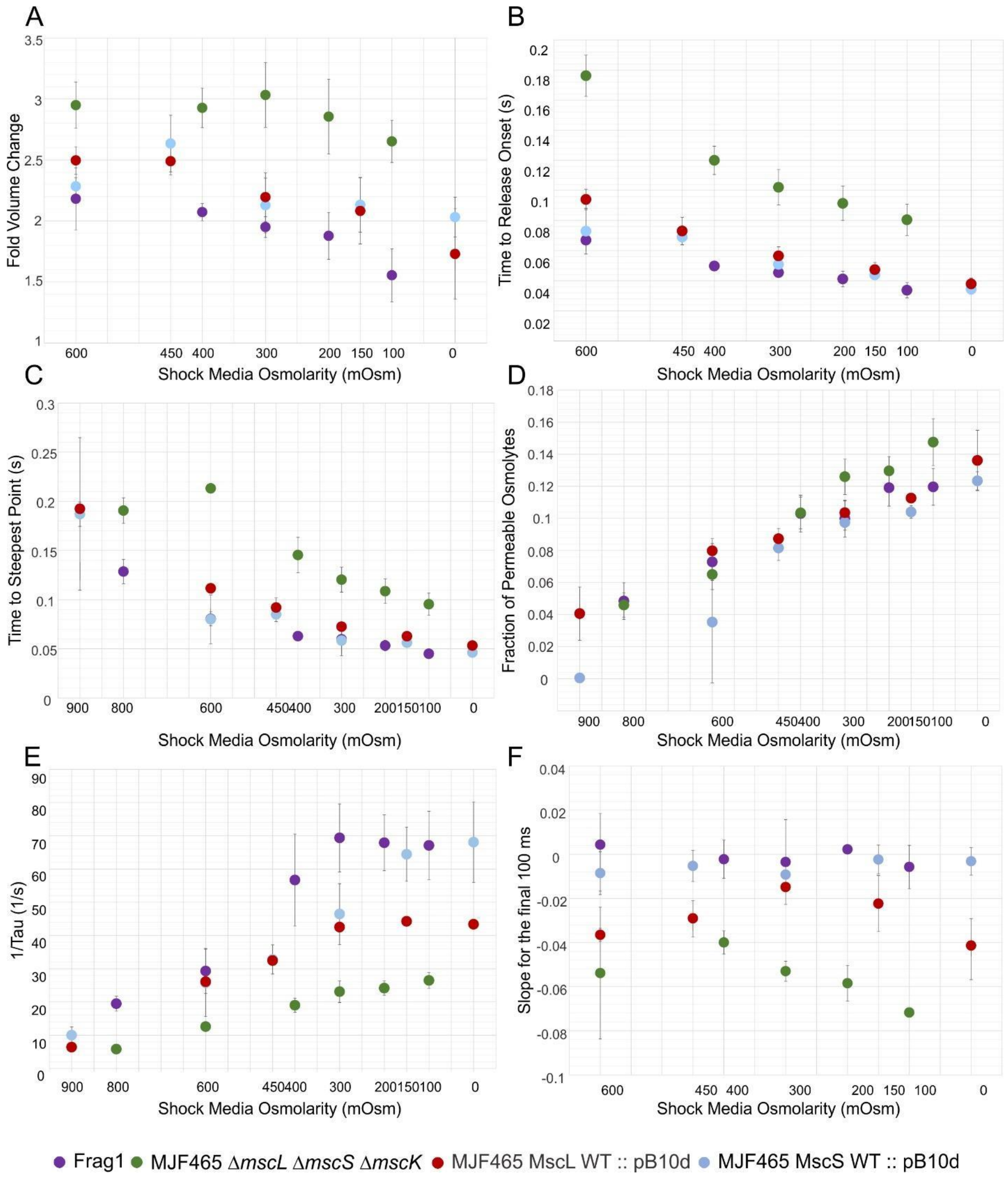


Supplemental Figure S7. Kinetic parameters of osmolyte release from five strains extracted from stopped-flow traces. The major difference is a higher negative slope of MJF465-MscL traces (panel F) that correlates with lower osmotic survival.

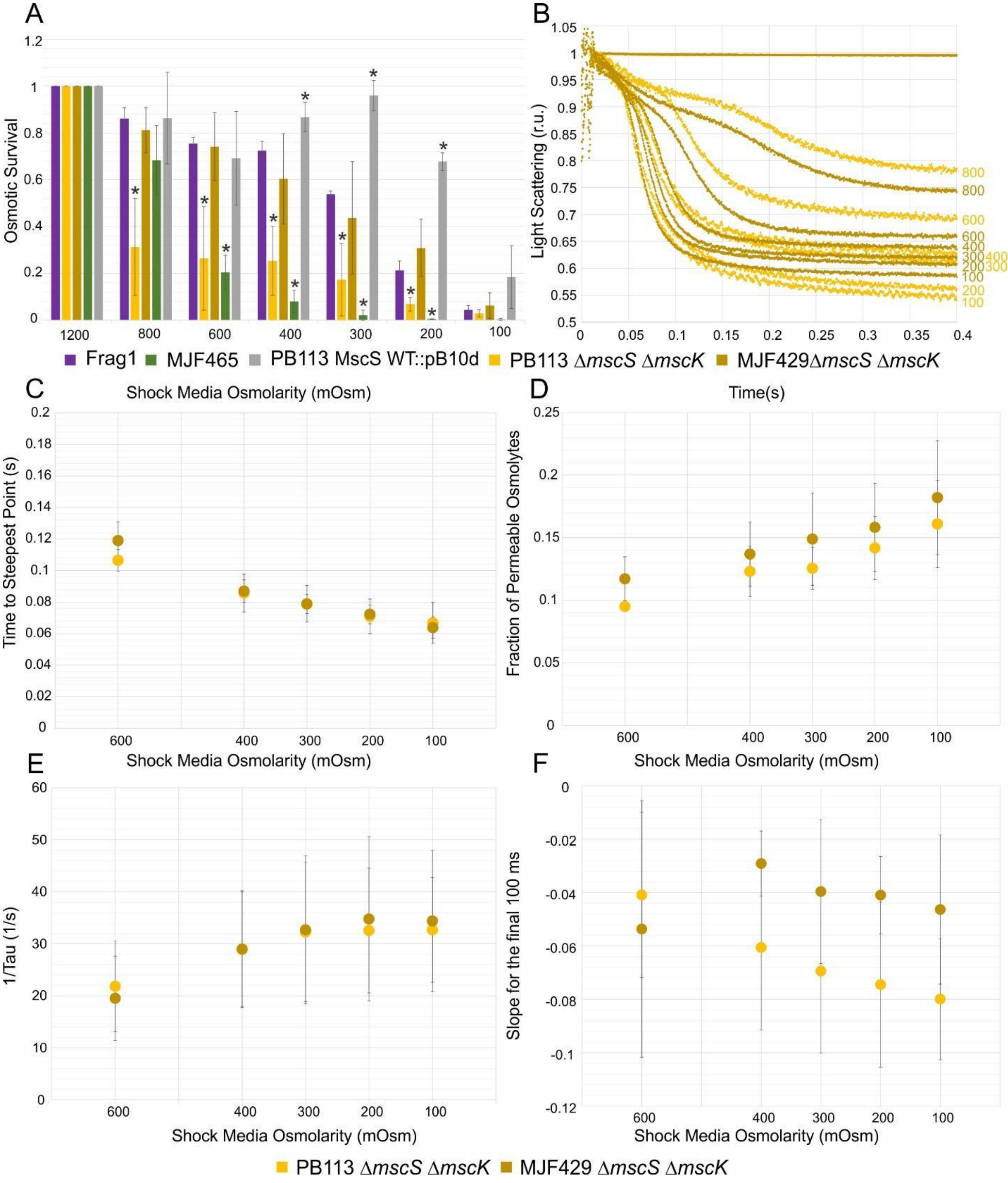


Supplemental Figure S8. The stopped-flow comparison of two *mscL*-carrying strains, MJF429 *(ΔyggB, ΔmscK*) and PB113 *(ΔyggB, ΔmscK, ΔrecA*). (A) Osmotic survival of the two strains in comparison with the positive (Frag1) and negative (MJF465) controls. Additional data show the rescuing of MJF465 and PB113 by introduction of WT MscS on a plasmid. Statistically significant ($p<0.05$) differences in survival values when compared to the Frag1 strain are represented with an asterisk. The full table of all p-values can be found in Supplemental Table S1. (B) The kinetics of light scattering for the two strains illustrates the major difference between the strains in the slope at the end of 0.4 s traces. Parameters extracted from these traces: time to steepest point (C), fraction of permeable osmolytes (D), the rate of release, 1/Tau, (E) and the slopes at the end of the trace (F). The data represent averages of multiple independent trials.

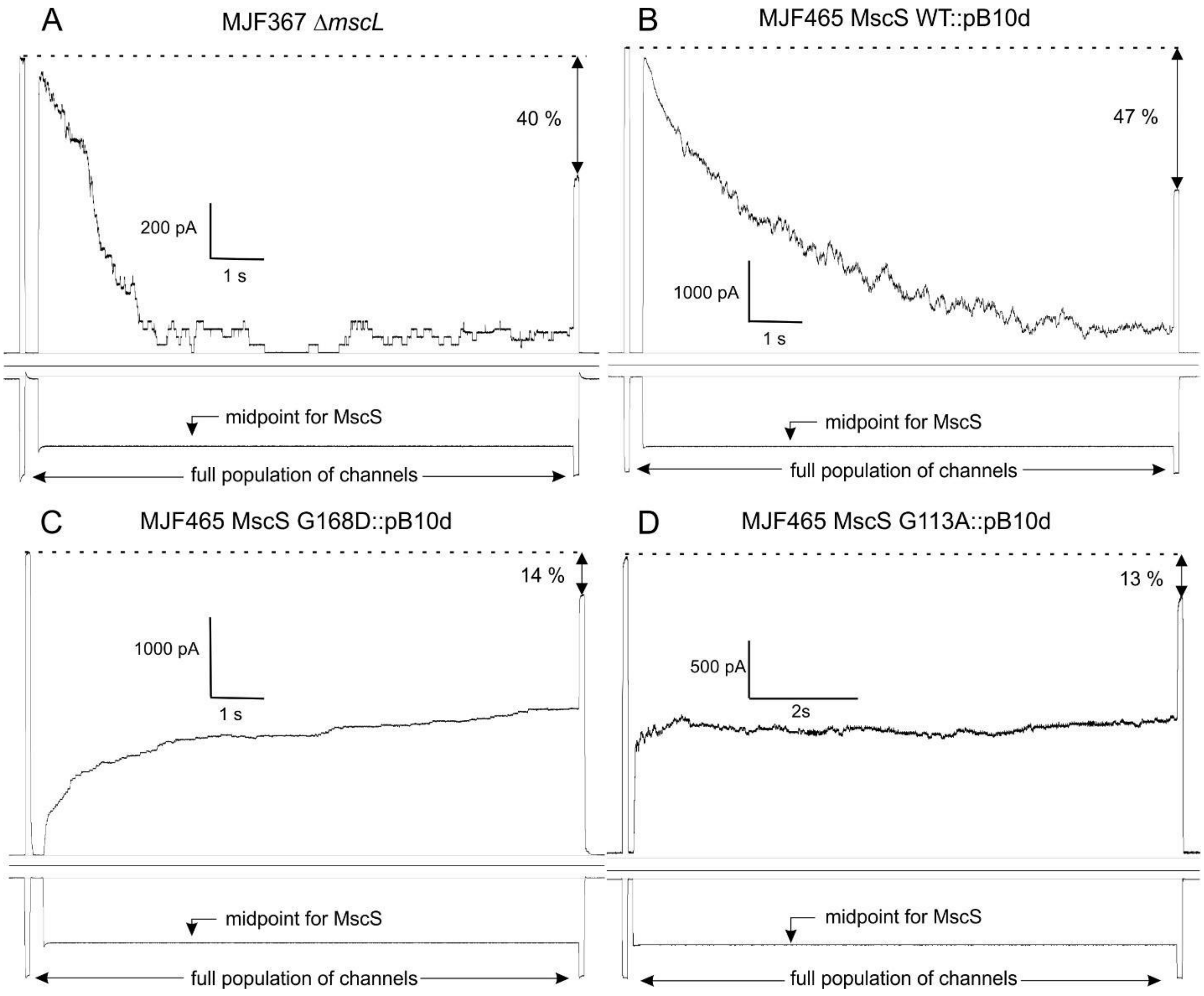


Supplemental Figure S9. Patch-clamp traces demonstrating inactivation for the native MscS (A), the WT MscS with plasmid expression (B), the G168D mutant (C), and the G113A mutant (D). The pulse-step-pulse protocol revealed the entire active channel population with the first saturating pulse. The following prolonged step of moderate pressure provided a conditioning period during which the channels had a chance to inactivate. The final saturating pressure pulse reveals the non-inactivated part of the channel population.

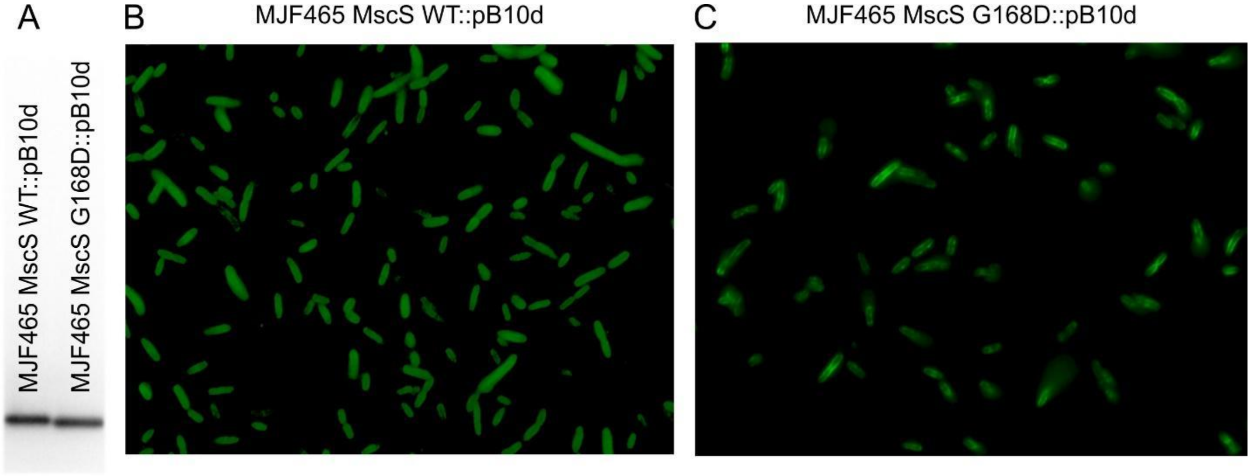


Supplemental Figure S10. Anti-6-His Western Blot and immunostaining comparing the expression of WT MscS with the G168D mutant.

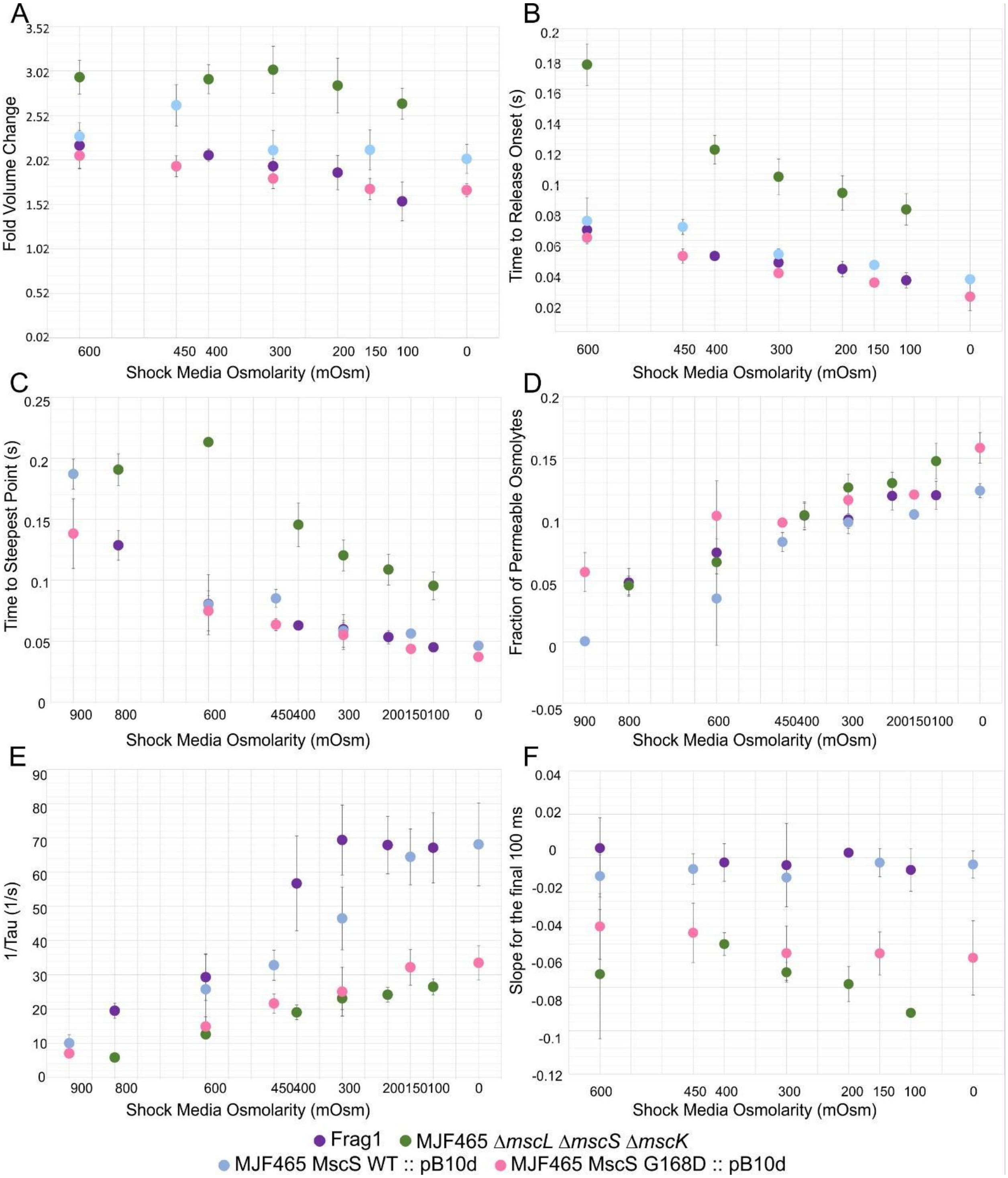


Supplemental Figure S11. Kinetic parameters of osmolyte release from stopped flow traces comparing the MscS G168D mutant and WT MscS with Frag1 and MJF465. The major differences between the traces are a longer Tau (panel E) and higher negative slope (panel F) of MscS G168D expressed in MJF465 that correlates with lower osmotic survival.

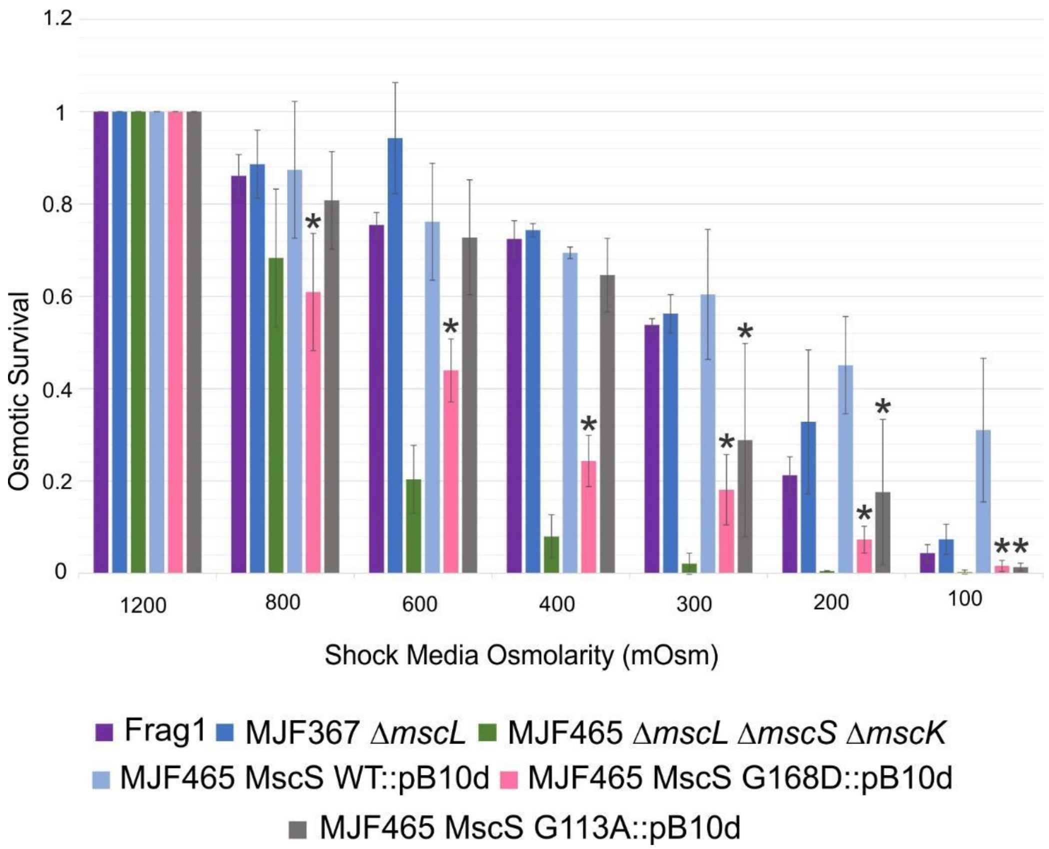


Supplemental Figure S12. Osmotic survival data comparing WT MscS with non-inactivating mutants expressed in the MJF465 triple knock-out strain. Data for MJF367 carrying the chromosomal copy of *mscS* and expressing the channel at its native level is included. Statistically significant ($p<0.05$) survival values of MscS mutations compared to plasmid expressed WT MscS are represented with an asterisk. The full table of all p-values can be found in Supplemental Table S1.

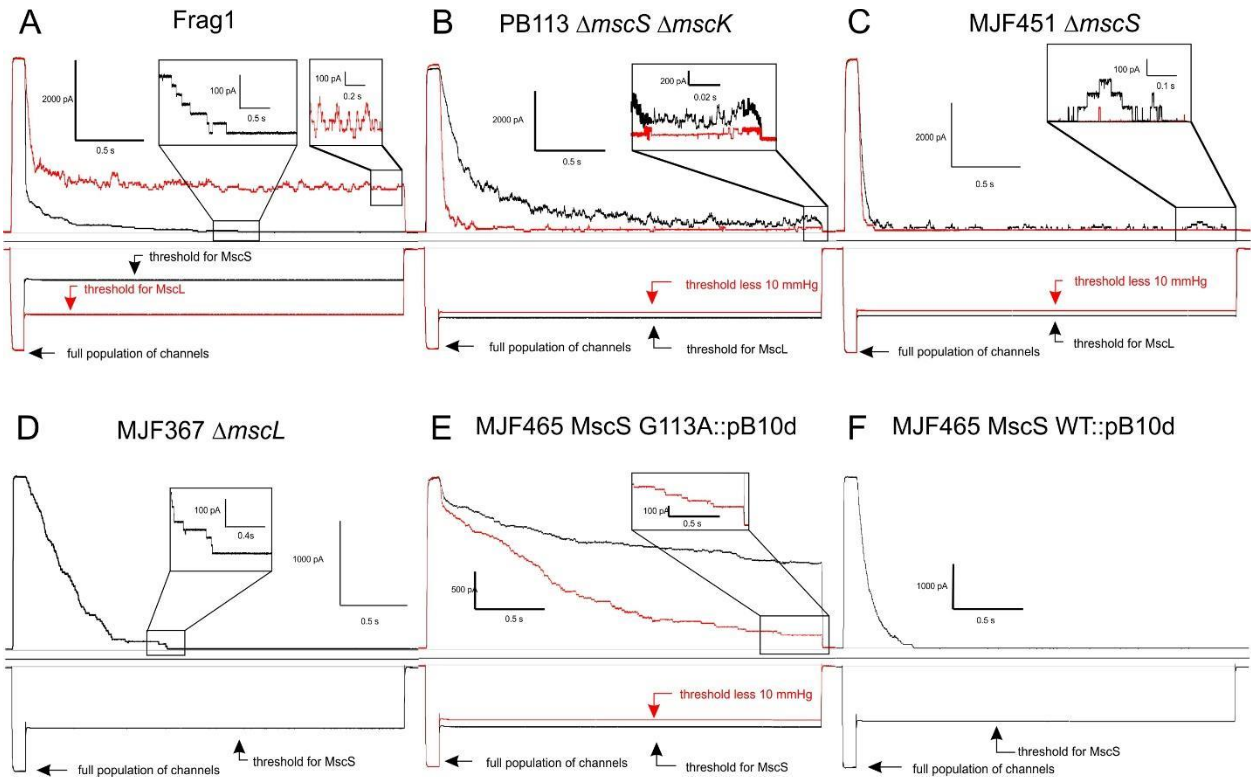


Supplemental Figure S13. Patch-clamp traces comparing Frag1, MJF451, and MJF367 using the same pulse-step protocol as in Figure 9. Frag1 (A) at the threshold for MscL (red) shows residual channel activity (right box) and at the threshold for MscS (black) shows termination of the response (left box). MJF451 with MscK (C) shows a reduced flickering at the threshold for MscL (black) and hardly any channel activity at 10 mmHg below the threshold (red). MJF367 (D) shows early and complete silencing of channel activity. (E) The G113A mutant expressed in MJF465 cells shows slow adaptation and lingering activity at the threshold tension. (F) WT MscS closes faster and completely at its threshold tension.
